# Dormancy-inducing 3D-engineered matrix uncovers mechanosensitive and drug protective FHL2-p21 signaling axis

**DOI:** 10.1101/2023.01.25.525382

**Authors:** Sadra Bakhshandeh, Unai Heras, Hubert M. Taïeb, Adithi R. Varadarajan, Susanna M. Lissek, Sarah M. Hücker, Xin Lu, Daniela S. Garske, Sarah A. E. Young, Andrea Abaurrea, Maria M Caffarel, Ana Riestra, Paloma Bragado, Jörg Contzen, Manfred Gossen, Stefan Kirsch, Jens Warfsmann, Kamran Honarnejad, Christoph A. Klein, Amaia Cipitria

**Author notes:** Both authors contributed equally to this manuscript (Sadra Bakhshandeh, Unai Heras).

## Abstract

Resected tumors frequently relapse with distant metastasis, despite systemic treatment. Cellular dormancy has been identified as an important mechanism underlying such drug resistance enabling late relapse. Nonetheless, hurdles associated with detection and isolation of disseminated cancer cells (DCCs) in disease-free patients urge the need for *in vitro* models of dormant cells suited for drug discovery. Here, we explore dormancy-inducing 3D-engineered matrices, which generate mechanical confinement and induce growth arrest and survival against chemotherapy in cancer cells. We characterized the dormant phenotype of solitary cells by P-ERK^low^:P-p38^high^ dormancy signaling ratio, along with Ki67-expression. As underlying mechanism, we identified stiffness-dependent nuclear localization of the four-and-a-half LIM domains 2 (FHL2) protein, leading to p53-independent high p21^Cip1/Waf1^ nuclear expression, validated in murine and human tissue. Suggestive of a resistance-causing role, cells in the dormancy-inducing matrix became sensitive against chemotherapy upon FHL2 downregulation. Thus, our biomaterial-based approach will enable systematic screens for novel compounds suited to eradicate potentially relapsing, dormant cancer cells.

**Teaser:** Using semi-synthetic bioengineered hydrogels, we reveal a mechanosensitive and drug protective mechanism of dormant cancer cells in tissues

## Introduction

Most cancer-associated deaths are due to metastasis, with eventual relapses spanning from years to decades despite adjuvant therapies (*1*, *2*). Estrogen receptor negative (ER-) breast cancer patients are often diagnosed with metastasis or relapse within the first 5 years, whereas patients with ER+ breast cancer continue to relapse beyond 8 years after surgery (*1*). Cancer dormancy has been described as one of the main mechanisms of evasion underlying late relapse (*1*, *3*). Due to its asymptomatic nature, it poses significant technical and ethical challenges for the identification, isolation and investigation of disseminated cancer cells (DCCs) in patients (*1*, *4*).

In solid epithelial cancer, detection of DCCs in asymptomatic patients without metastases (M0) usually relies on antibodies against proliferation markers such as Ki67. These markers, defined decades ago, are insufficient to characterize DCC populations due to their weak correlation to differentiate quiescent and dormant states, and their functional phenotypic variations, which complicate the detection of early and late DCCs (*1*, *5*). In current clinical procedures, solitary DCCs are too small to be detected by imaging technologies, since the inaccessibility to specific niches remains a barrier for routine procedures (*5*). All these constraints hinder the understanding of the survival and escape mechanisms of DCCs, which differ from proliferative tumor masses. Therefore, gaining new insights into their nature, functional phenotypes, proteomics signature and interaction with the microenvironment and extracellular matrix (ECM) is critical for future diagnostic DCC detection and targeting for drug discovery.

Recent *in vivo* dormancy models have further revealed the role of the ECM on breast cancer cell (BCC) dormancy, as highlighted with the effect of the architecture of tumor-derived collagen type III (*6*), astrocyte-deposited laminin-211 (*7*), adhesive glycoprotein thrombospondin-1 (*8*), integrin-mediated interactions (*9*, *10*) or neutrophil extracellular traps (*11*). Despite all the progress made, the availability of robust animal models to study dormancy is still limited. Current *in vivo* models do not typically consider all the phenotypic variation of DCCs from non-metastatic to fully metastatic states (*1*, *3*, *6*, *8*, *10*–*13*). Moreover, characterization of cellular dormancy or single DCCs remains extremely challenging due to their low abundance (*5*). All these limitations call for reliable *in vitro* culture systems to mimic niche-specific microenvironmental architectures and genotypic/phenotypic states of single DCCs, to finally enable scalable drug screening campaigns for the identification of promising DCC-selective drugs and drug targets.

Overcoming the limits of Matrigel as a standard *in vitro* model for drug screening (*14*, *15*), semi-synthetic biomaterial-based approaches have been previously pursued to generate bioengineered niches mimicking tissue-specific ECM. Biomaterial-based matrices have been shown to induce and/or preserve quiescence of astrocytes (*16*) and muscle stem cells (*17*), maintain neural progenitor (*18*) and hematopoietic cell stemness (*19*), investigate mechanical memory (*20*), differentiation of stem cells (*21*), as well as cancer cell cycle progression (*22*, *23*).

These systems have incredible potential to analyze stromal and parenchymal ECM contributions through simple interactions with cells, or even hypoxia and matrix mechanics to mimic cellular and tumor dormancy (*24*). In particular, *in vitro* models of breast cancer dormancy harness specific biophysical and biochemical environmental cues to influence cellular behavior through controlled cell-environment interactions. Tunable biophysical properties such as stiffness or viscoelasticity in versatile hydrogels have enabled exploring proliferative, growth-arrested or dormant phenotypes of cancer cells, largely via 3D physical confinement (*25*–*28*). Stiffer gels have been associated with more dormant phenotypes, whereas fast stress-relaxation has been linked to a transition from growth arrest to proliferation (*23*). Interestingly, biochemical properties such as integrin binding and matrix metalloproteinase (MMP) degradation profiles can also recapitulate dormancy induction at metastatic sites (*24*). Poly(ethylene glycol) hydrogels manipulated with MMP-cleavable crosslinkers and arginine-glycine-aspartic acid (RGD) integrin-binding sites have uncovered mechanisms underlying dormancy induction and reactivation of BCCs (*22*, *29*, *30*). As for other biochemical properties, such as nutrient deprivation in growth arrested cells, serum starvation has been used to explore the reorganization of fibronectin secreted by dormant cells (*31*). This reorganization was necessary for the survival of dormant cancer cells, while oxidation or proteolytic degradation of ECM proteins could shift the effect towards proliferative and invasive phenotypes (*31*, *32*). Serum starvation was also studied in 3D collagen and Matrigel, where Matrigel supported a large population of dormant D2.0R cells and collagen contained more cells with senescent behavior (*33*).

Notably, most of these models have been capable of recapitulating tumor mass dormancy, whereby tumor masses are held in balance, with cell proliferation rate equal to cell death. As a result, some of these dormancy-inducing models have been tested with potential drugs (*14*, *25*, *34*– *36*). Although tumor mass dormancy is one category of dormancy, it does not fully capture the functional phenotype of cellular dormancy of solitary DCCs. Therefore, high-throughput drug screening campaigns to target solitary dormant DCCs in bioengineered niches are still scarce (*24*).

Here we harness matrix stiffness to induce a dormant phenotype of solitary cancer cells within a few days, offering new opportunities for potential high-throughput drug screening of single dormant DCCs. We explored ultraviolet (UV) light-initiated, thiol-ene-mediated, covalently-crosslinked inert alginate hydrogels, which primarily through 3D mechanical confinement can induce and maintain large populations of single BCCs in a dormant state. As extensively used in previous *in vitro* models (*23*–*26*, *28*), here we selected commonly-used BCCs, MDA-MB-231 as a model for triple-negative breast cancer and MCF-7 for the luminal subtype. Consciously using non-adhesive and non-degradable alginate hydrogels, we show a majority of surviving single cells in the G_0_/G_1_ cell cycle phase, based on the fluorescence-ubiquitination-based cell cycle indicator 2 (FUCCI2). We demonstrate that the survival of these cells is mainly dominated by matrix stiffness, capable of inducing a dormancy signature through the well-established P-ERK^low^:P-p38^high^ ratio (*37*, *38*), along with a majority of Ki67 negative (Ki67-) cells. Combined with RNA sequencing, we revealed the stiffness-dependent nuclear localization of the four-and-a-half LIM domains 2 (FHL2) protein as an underlying mechanism of dormant DCCs in 3D alginate gels, leading to a p53-independent high p21^Cip1/Waf1^ nuclear expression, validated in murine and human tissue. Notably, we observed that this effect endowed growth-arrested cells with resistance to cell cycle-specific chemotherapy, such as paclitaxel, mainly effective in the S/G_2_/M phase. Suggestive of a growth arrest-mediated drug resistance, dormant DCCs became sensitive against drug treatment upon FHL2 downregulation. This 3D biomaterial-based approach with mechanically-induced dormant single cancer cells offers a simple and scalable method. It enables investigating large populations of dormant single cancer cells in a few days, which are otherwise rare and inaccessible in large numbers from a clinical setting, and could pave the path towards therapeutic applications, e.g., via high throughput drug screens of novel compounds to eradicate potentially relapsing DCCs.

## Results

### 3D mechanical confinement via covalently-crosslinked alginate yields distinct fractions and populations of dormant cells based on hydrogel composition

Building on the now widely acknowledged role of the ECM in cancer induction and progression, we employed a versatile ECM-mimicking hydrogel encapsulation approach to investigate the effect of ECM-mediated 3D mechanical confinement on cell cycle progression. We used ultraviolet (UV) light-initiated thiol-ene-crosslinked alginate (*39*), an inherently inert hydrogel (non-cell adherent and non-degradable), which allows for selective modulation of parameters such as stiffness, cell adhesion or degradation (Fig.1a). Cells were encapsulated in the presence (or absence) of adhesion ligands (RGD), within degradable MMP-sensitive or non-degradable dithiothreitol (DTT)-crosslinked hydrogels (Fig.1a and Supp.Fig.S1). The stiffness of the hydrogels ranged between 0.5-13 kPa as a function of crosslinker concentration (Fig.1b). To monitor cell cycle progression in real-time, we genetically modified widely-used triple-negative MDA-MB-231 and luminal MCF7 BCC lines, known to disseminate to secondary organs in *in vivo* models, with the fluorescence-ubiquitination-based cell cycle indicator 2 (FUCCI2) (*40*, *41*) (Fig.1c). Time-lapse imaging of MDA-MB-231-FUCCI2 cells encapsulated within alginate hydrogels of different stiffness (soft for 1 kPa and stiff for 10 kPa), adhesion and degradation properties, as well as on two-dimensional tissue culture plastic (2D-TCP) and in 3D Matrigel, was performed over a period of 4-5 days, followed by quantification of the cell cycle state at the experimental endpoint (Fig.1d, Supp.Mov.1). 2D TCP and 3D Matrigel yielded highly proliferative (Ki67+) and metabolically active cells (resazurin reduction) in the form of single cells and 3D spheroids, respectively (Fig.1d, Supp.Fig.S2b,d). Cells within the 3D alginate groups remained mostly as single cells, with the highest fraction of S/G_2_/M phase cells being in the soft group, while the stiff adherent (RGD) and degradable (MMP-sensitive) groups displayed a lower percentage of S/G_2_/M and higher fraction of G_0_/G_1_ phase cells (Fig.1d). 3D alginate stiff yielded the highest fraction of cell cycle-arrested cells over 5 days (Fig.1d), while preserving high viability (Supp.Fig.S2a). Interestingly, we noticed different sub-populations of cell cycle-arrested cells as a function of the expression (or lack of) of the licensing factor cdt1 (G_0_/G_1_ cdt1+/–; Fig.1d, Fig.2a, Supp.Mov.2-7). A recent mapping of protein dynamics during cell cycle progression revealed that the expression patterns of cdt1 are highly dynamic and distinctive of different levels of quiescence, with the lowest (cdt1-) identifying a longer time spent in a growth-arrested state (*42*).

**Fig. 1.**
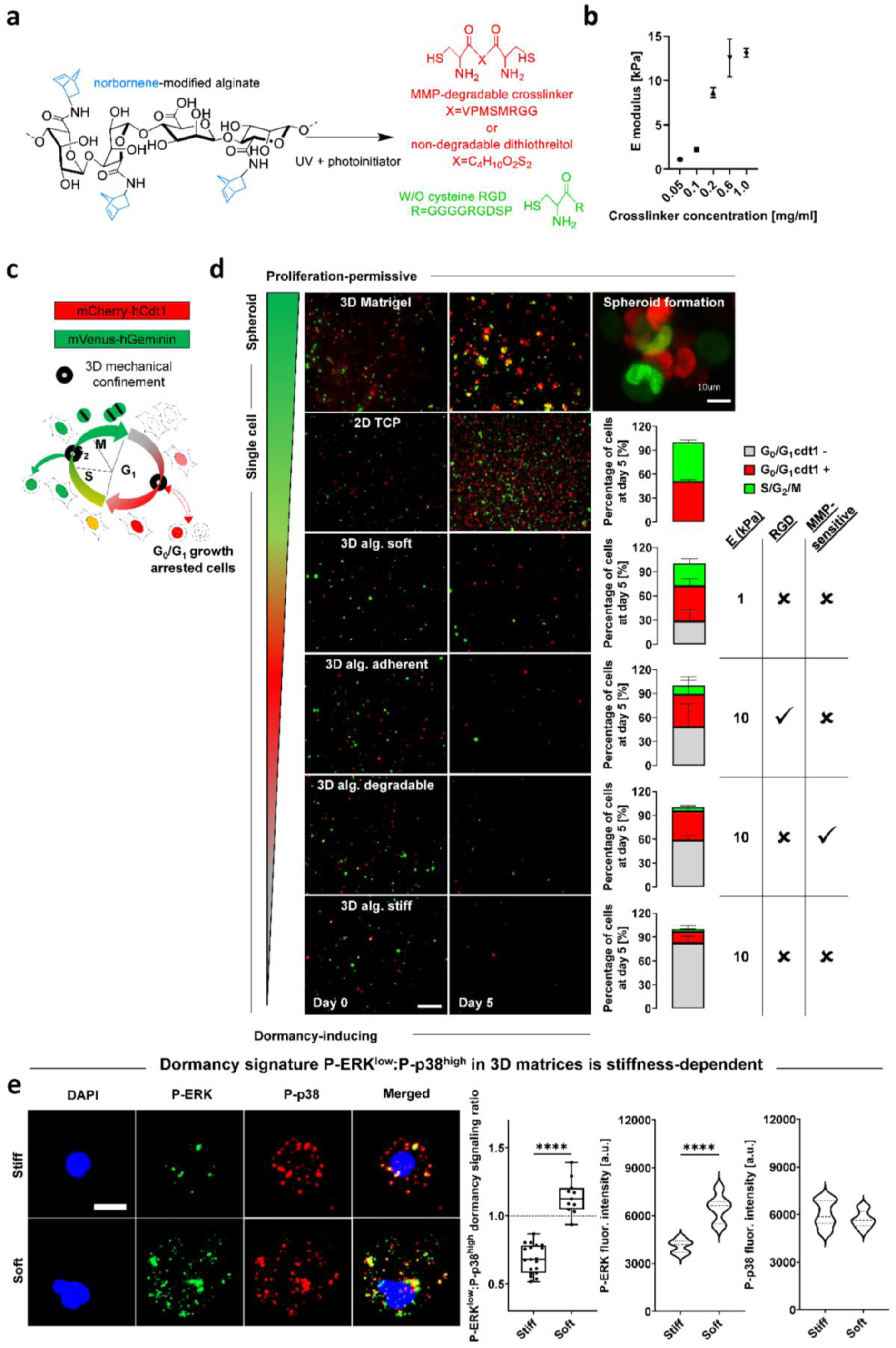
3D mechanical confinement via covalently-crosslinked alginate yields distinct fractions and populations of growth-arrested cells based on hydrogel composition. **a)** Bis-cysteine enzymatically (MMP) degradable peptide or non-degradable dithiothreitol (DTT) (red) crosslink with norbornene-modified alginate (blue-black) in the presence or absence of cysteine-coupled RGD molecules (green). **b)** Frequency sweep of norbornene-modified alginate (no RGD) with different concentrations of DTT crosslinker yields a range of stiffness (Elastic/Young’s moduli) (n=3). **c)** Diagram of FUCCI2 cell cycle reporter activity (G_0_/G_1_=mCherry-hCdt1 and S/G_2_/M=mVenus-hGeminin). The cartoon was adapted from Sakaue-Sawano et al. (*13*), Copyright (2008), with permission from Elsevier. **d)** Time lapse imaging (5 days) of MDA-MB-231-FUCCI2 cells within a range of proliferation-permissive and dormancy-inducing hydrogels with distinct stiffness, adhesion and degradation properties (right-hand side table). Representative time lapse fluorescence maximum projections at day 0 and day 5, scale bar equals 200 μm. Scale bar equals 10 μm for 3D Matrigel spheroid image. Percentage bar plots show fraction of cell cycle distribution at day 5 for different experimental groups (n≥3 gels for 82 to 453 cells). **e)** Representative images of P-ERK and P-p38 for MDA-MB-231 cells within 3D stiff and soft alginate hydrogels after 5 days of encapsulation (blue=DAPI, green=P-ERK, red=P-p38). Scale bar equals 10 μm, P-ERK^low^:P-p38^high^ ratio quantification in based on each phosphorylated protein fluorescence intensity (a.u.=arbitrary unit) per cell (n=10-20 single cells per condition). Student’s t-test, ****p<0.0001.

**Fig. 2.**
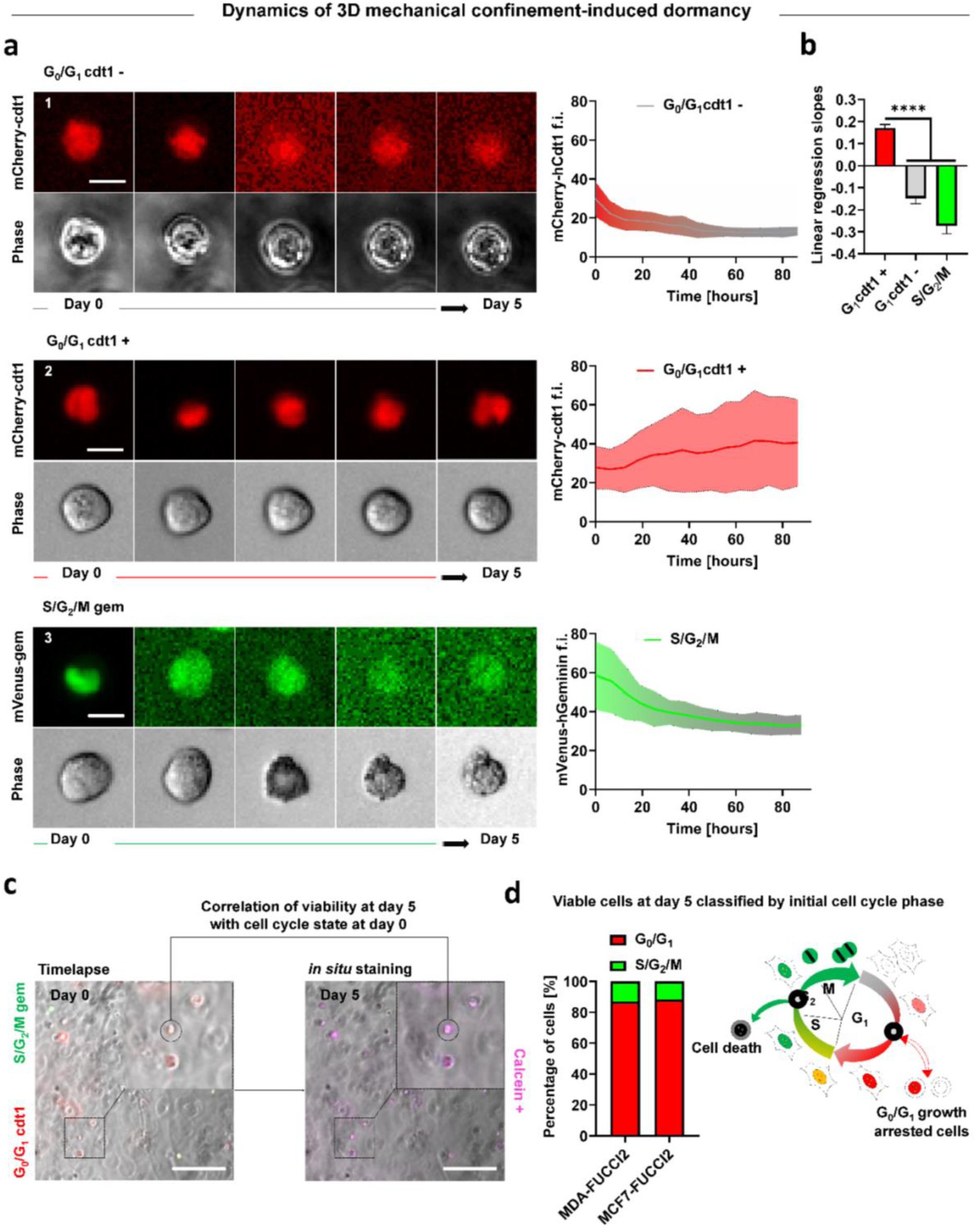
Dynamics of cell cycle and viability correlation reveal that G_0_/G_1_ cells are substantially more resistant to 3D mechanical confinement than cells in the S/G_2_/M phase. **a)** Representative time lapse images of single MDA-MB-231-FUCCI2 cells in 3D alginate stiff hydrogels with respective single cell longitudinal tracking of mCherry-hCdt1 and mVenus-hGeminin fluorescence intensity (f.i.). Center trace and shaded area indicate mean and standard deviation (n≥15 cells). Scale bar equals 10 μm. **b)** Comparison of linear regression slopes from f.i. tracking graph. Kruskal–Wallis test with Dunn’s correction, ****p<0.0001. **c)** Experimental workflow for longitudinal correlation of FUCCI2 initial cell cycle state within 3D alginate stiff hydrogels and viability after 5 days of encapsulation. Cells were stained *in situ* for viability after 5 days in time lapse and the selected cells (calcein+) were tracked back to their initial cell cycle state (time of encapsulation). Scale bar equals 200 μm. **d)** Percentage bar plots show fraction of viable cells with respect to their initial cell cycle state for MDA-MB-231-FUCCI2 and MCF7-FUCCI2 cells (n≥241 cells pooled from 3 independent experiments). Diagram of cell cycle progression under 3D mechanical confinement, revealing growth arrest and higher resilience for cells in G_0_/G_1_ and death for cells in S/G_2_/M. The cartoon was adapted from Sakaue-Sawano et al.^40^, Copyright (2008), with permission from Elsevier.

We further explored this stiffness-induced growth arrest state by the well-established ratio of phosphorylated forms P-ERK^low^:P-p38^high^ as a dormancy marker (*3*, *29*, *33*, *37*, *38*). We analyzed the ratio at a single cell level within 3D stiff and soft alginate hydrogels using immunofluorescence (Supp.Fig.S2e). We determined a significant decrease in P-ERK proliferative activity of cells in stiff alginate gels compared to soft gels (Fig.1e, Supp.Fig.S2e). Expression of P-p38 was heterogeneous in both types of matrices (Fig.1e, Supp.Fig.S2e). Therefore, we confirmed a significantly lower P-ERK^low^:P-p38^high^ ratio as a maker for dormancy of single cells embedded in stiff alginate gels. Finally, reminiscent of post-therapeutic relapses in clinical settings, growth-arrested cells were able to exit this dormant state and resumed proliferation when retrieved from the hydrogels and seeded on 2D TCP (Supp.Fig.S2c).

### Dynamics of cell cycle and viability correlation reveal that G_0_/G_1_ cells are substantially more resistant to 3D mechanical confinement than cells in the S/G_2_/M phase

To understand the dynamics of 3D mechanical confinement-induced growth arrest, we tracked single-cell MDA-MB-231-FUCCI2 fluorescence intensity (f.i.) within stiff hydrogels. Cells which were in the G_0_/G_1_ state at the time of encapsulation had either lost (cdt1-) (Fig.2a1, b) or kept (cdt1+) (Fig.2a2, b) their fluorescence by the end of the experiment. On the other hand, in almost every cell initially in the S/G_2_/M phase, we observed a decrease in f.i. within the first few hours after encapsulation (Fig.2a3, b). By correlating cell’s FUCCI2 imaging and *in situ* viability staining (calcein+; Fig.2c), we revealed that the vast majority of viable cells at day 5 were initially in the G_0_/G_1_ cycle state (Fig.2c, d). We validated these results for MCF7-FUCCI2 cells (Fig.2d), confirming that cells in the G_0_/G_1_ state are more resilient to 3D mechanical confinement compared to cells in the S/G_2_/M phase.

### Growth-arrested viable BCCs under 3D mechanical confinement are associated with increased chemoresistance to cell cycle-specific drugs

Chemoresistance of growth-arrested triple negative MDA-MB-231 cells, upon 3D mechanical confinement in 3D alginate stiff hydrogels, was compared with the commonly-used proliferation-permissive 3D Matrigel, and tested with four different cytotoxic drugs. We used paclitaxel, 5-fluorouracil and gemcitabine as standard cell cycle-specific drugs used in adjuvant therapy for triple negative breast cancer patients (*43*, *44*). We further tested the effect of carboplatin, a cell cycle non-specific alkylating agent. The main readouts were fold change in cell viability and metabolic activity with respect to vehicle.

Considering 0.01 mM of paclitaxel a standard drug dose, at lower doses of paclitaxel (≤0.01 mM) we observed significant decrease in viability of proliferating cells in 3D Matrigel, but not for growth-arrested MDA-MB-231 cells in 3D alginate stiff (Fig.3a,b,c). At higher doses (>0.01 mM) the viability dropped for both cells in Matrigel and alginate stiff. This effect of chemoresistance was further enhanced with the fold change in metabolic activity, where there was a significant decrease for proliferating cells in Matrigel but not for growth-arrested cells in alginate stiff, at both low and high doses of paclitaxel (Fig.3d).

**Fig. 3.**
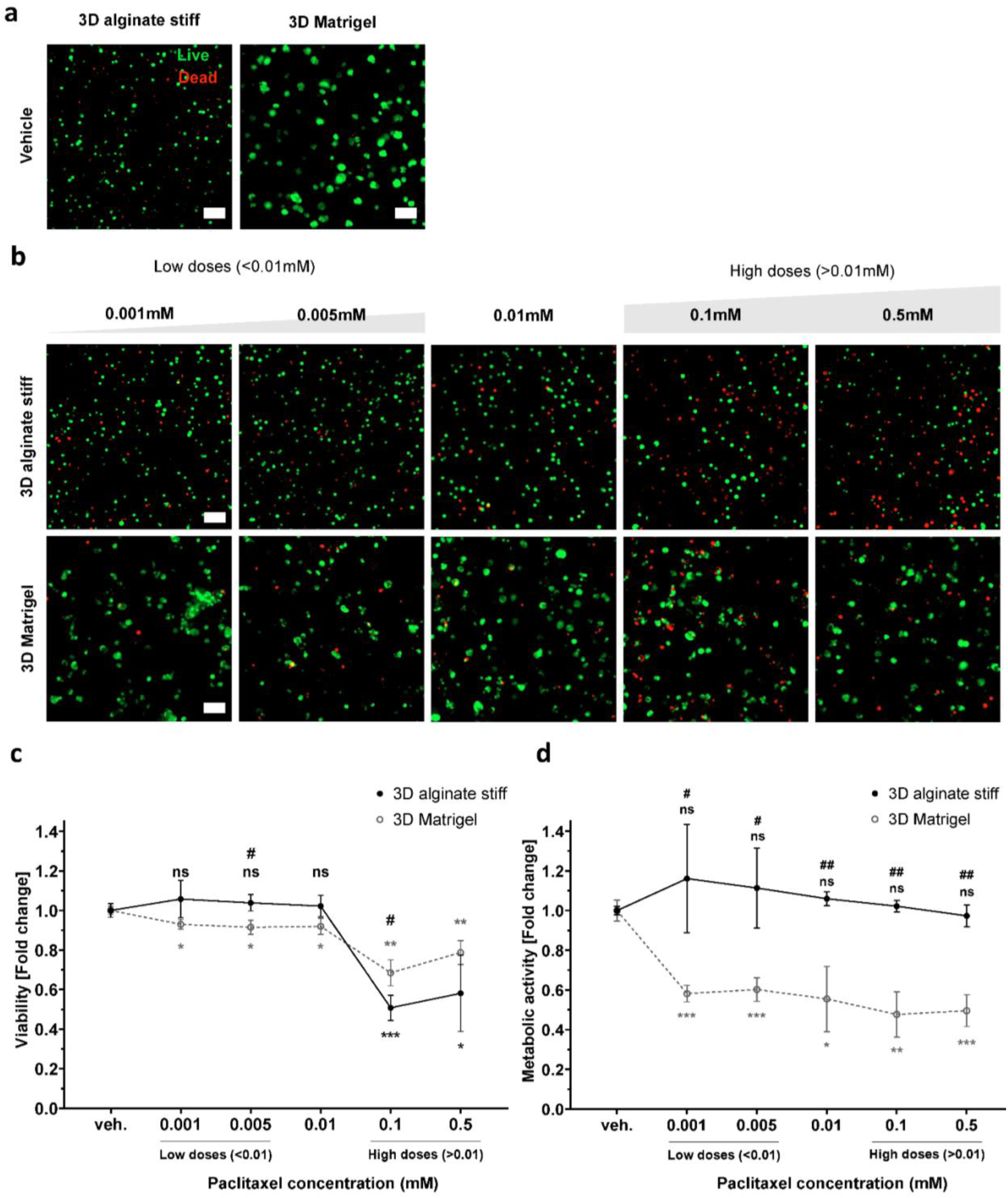
Growth-arrested viable BCCs within 3D alginate stiff gels are more resistant to cell cycle-specific paclitaxel. **a)** Representative live/dead (live=calcein, dead=ethidium homodimer) maximum projection fluorescence images of MDA-MB-231 cells encapsulated in dormancy-inducing 3D alginate stiff and proliferation-permissive 3D Matrigel vehicles for 5 days without drug exposure (n=3 gels with 150 to 250 cells). Scale bar equals 100 µm. **b)** Representative live/dead (live=calcein, dead=ethidium homodimer) maximum projection fluorescence images of MDA-MB-231 cells encapsulated in dormancy-inducing 3D alginate stiff and proliferation-permissive 3D Matrigel for 3 days and exposed to paclitaxel (0.001 to 0.5 mM) for 2 days (n=3 gels with 150 to 250 cells). Scale bar equals 100 µm. **c)** Quantification of viability fold change within 3D alginate stiff and 3D Matrigel with respect to the vehicle (untreated groups) and between groups. **d)** Quantification of metabolic activity fold change within 3D alginate stiff and 3D Matrigel with respect to the vehicle of each gel (untreated groups). * for statistical significance with respect to the vehicle of each gel (untreated groups) and # for statistics between groups. Student’s t-test (n=3 hydrogels), */#p≤0.05, **/##p≤0.01, ***/###p≤0.001.

Notably, proliferative MDA-MB-231 cells in 3D Matrigel showed high resistance to the other two cell cycle-specific drugs 5-fluorouracil and gemcitabine, even at concentrations up to 0.5mM (Supp.Fig.S3). We further tested the effect of carboplatin, a cell cycle non-specific alkylating agent. MDA-MB-231 cells also showed resistance to carboplatin in 3D Matrigel due to its ineffectiveness to BRCA1/2-deficient cells (*45*). For these reasons, we excluded these drugs for our analysis with growth-arrested cells in 3D alginate stiff. As a strategy, we aimed to identify an effective and sensitive axis to target dormant BCCs using cell cycle-specific paclitaxel.

### Gene expression analysis of cells in dormancy-inducing versus proliferation-permissive 3D matrices

We then looked into the gene expression of dormancy-induced cells in more detail. RNA-sequencing on live cells retrieved from 3D alginate stiff (Fig.4a) revealed enrichment of cell cycle and retinoblastoma-associated molecular pathways (Fig.4b-d, SuppFig.S4, 5), as well as regulation of senescence, autophagy and apoptosis processes when compared to cells retrieved from 3D Matrigel. In particular, we noticed strong enrichment of inflammatory response-associated pathways (TNF, NFkB and IL-17 signaling) and DNA damage stimuli response (ATM, p53 and miRNA regulation of DNA damage) (Fig.4b-e, Supp.Fig.S4, 5). Intriguingly, we noticed that despite increased mRNA expression of *cdkn1a* (coding for the p21 protein) upon culture in 3D alginate stiff compared to 3D Matrigel, the levels of its key upstream cell cycle regulator *p53* are lower (Fig.4e), hinting to a potential p53-independent mechanism regulating p21 expression (*46*). RNAseq results further revealed a dormant phenotype of non-proliferative MDA-MB-231 cells by the upregulation of two cyclin-dependent kinase (CDK) inhibitors p21 (*cdkn1a*) and p27 (*cdkn1b*) in 3D alginate stiff compared to 3D Matrigel (Supp.Fig.S6a). Reverse transcription quantitative PCR (RT-qPCR) analysis validated these results, further confirming the upregulation of CDK inhibitors p21 and p27 expression independent from p53 in 3D alginate stiff compared to 3D Matrigel (Supp.Fig.S6b). Furthermore, this matches the lack of correlation of p53-p21 observed in patients with estrogen receptor negative (ER-) breast cancer (Supp.Fig.S7). Noticing the particularly high fold change of p21 expression among other genes in 3D alginate (Fig.4e) and given its relevance in cell cycle regulation, we decided to investigate the role of p21 in regulating mechanically-induced cancer cell cycle arrest in more detail.

**Fig. 4.**
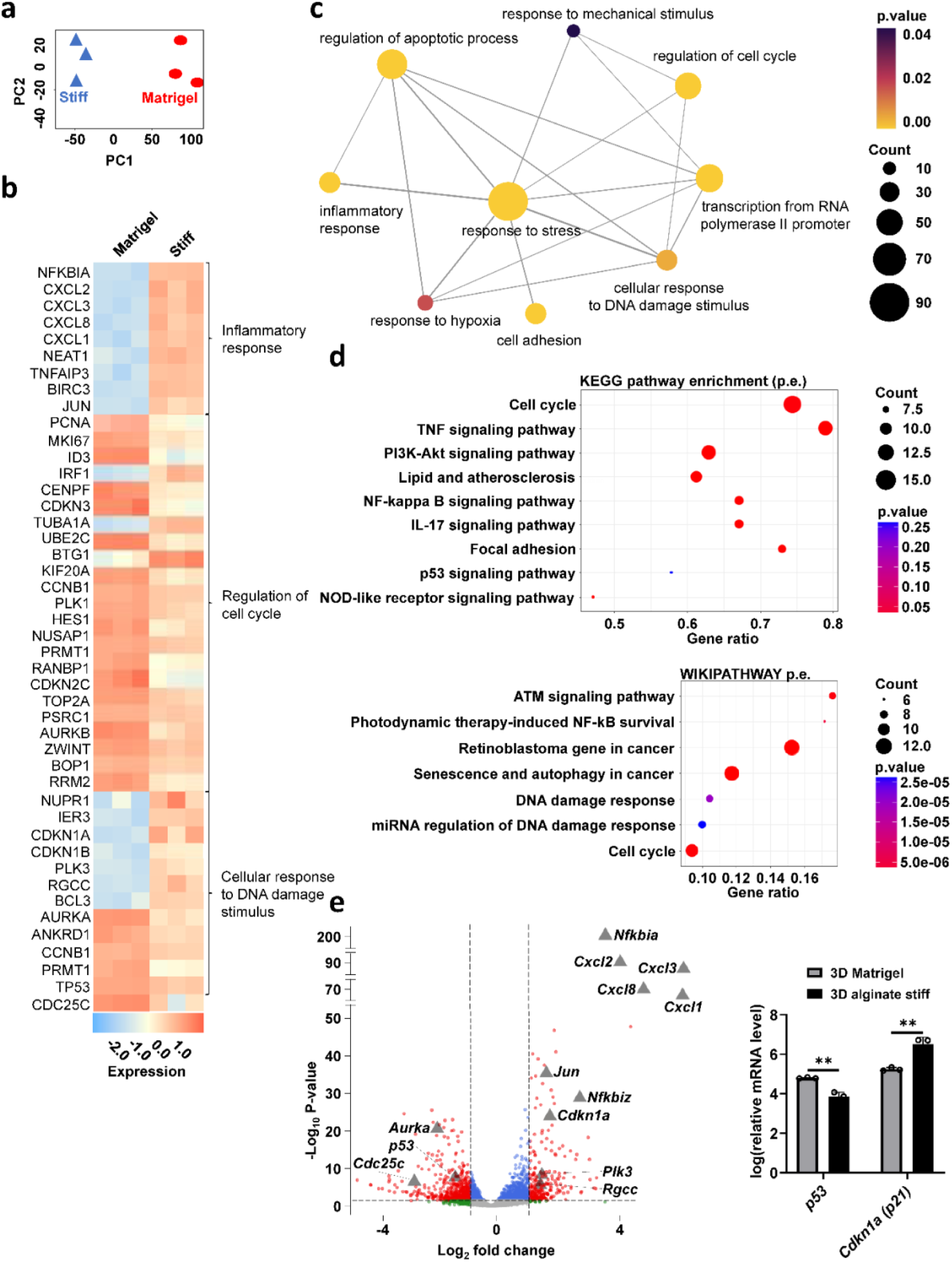
Gene expression analysis of cells in 3D alginate stiff compared to 3D Matrigel. **a)** Principal-component analysis (PCA) based on all expressed genes between 3D alginate stiff and 3D Matrigel groups acquired from RNA sequencing (n=3 for each condition). **b)** Heat map of selected differentially expressed genes involved in inflammatory response, regulation of cell cycle and cellular response to DNA damage stimulus. **c)** Network map of selected biological processes most enriched in 3D alginate compared to 3D Matrigel. Node size, node color and edge width represent number of genes, *p*-value from enrichment analysis and overlap of number of genes between two gene sets, respectively. **d)** Selected top differentially regulated pathways from Gene-set-enrichment analysis (GSEA) on differentially expressed genes between cells grown in 3D alginate stiff and 3D Matrigel according to KEGG and WIKIPATHWAY databases. **e)** Volcano plot of differentially expressed genes between 3D Matrigel and 3D alginate stiff. Grey triangles mark genes of interest associated with inflammation and DNA-damage pathways (grey dots=not significant, green dots= significant absolute log(base2) fold change of > 1, blue dots= significant *p-value* of < 0.05, red=significant absolute log(base2) fold change of > 1 and *p-value* <0.05 from a total of 9426 entries). Relative mRNA levels of p53 and *cdkn1a* (p21) (n=3 for each condition). Student’s *t*-test, **p≤0.01.

### p21 and FHL2 localization in 3D matrices is stiffness-dependent and FHL2 mediates p21 localization and Ki67 expression

Despite the now widely acknowledged advantages of 3D vs. 2D culture systems in studying physiologically-relevant cellular processes (*47*), the role of dimensionality in the expression, localization and function of p21 remains underexplored. This is of utmost importance considering that in cancer cells the p21 oncogenic vs. suppressing function is known to depend on its level of expression and intracellular localization (*46*). We therefore assessed these features across different 3D culture systems, focusing on soft and stiff 3D alginate hydrogels. As expected, the fraction of p21-positive cells was significantly higher in 3D alginate stiff compared to soft or 3D Matrigel (Supp.Fig.S8), accompanied by strong enrichment of DNA repair-associated pathways (Fig.5a, b and Supp.Fig.S9). Importantly, cells in 3D alginate stiff revealed a significantly higher nuclear vs. cytoplasmic (nuc./cyto. ratio) localization of p21 compared to the alginate soft group, together with a lower fraction of proliferative cells (Ki67 expression) (Fig.5c). Inhibiting MEK/ERK, PI3K/Akt and JNK, as some of the major signaling pathways regulating cell proliferation, displayed a strong effect on p21 localization (Fig.5d, Supp.Fig.S10).

**Fig. 5.**
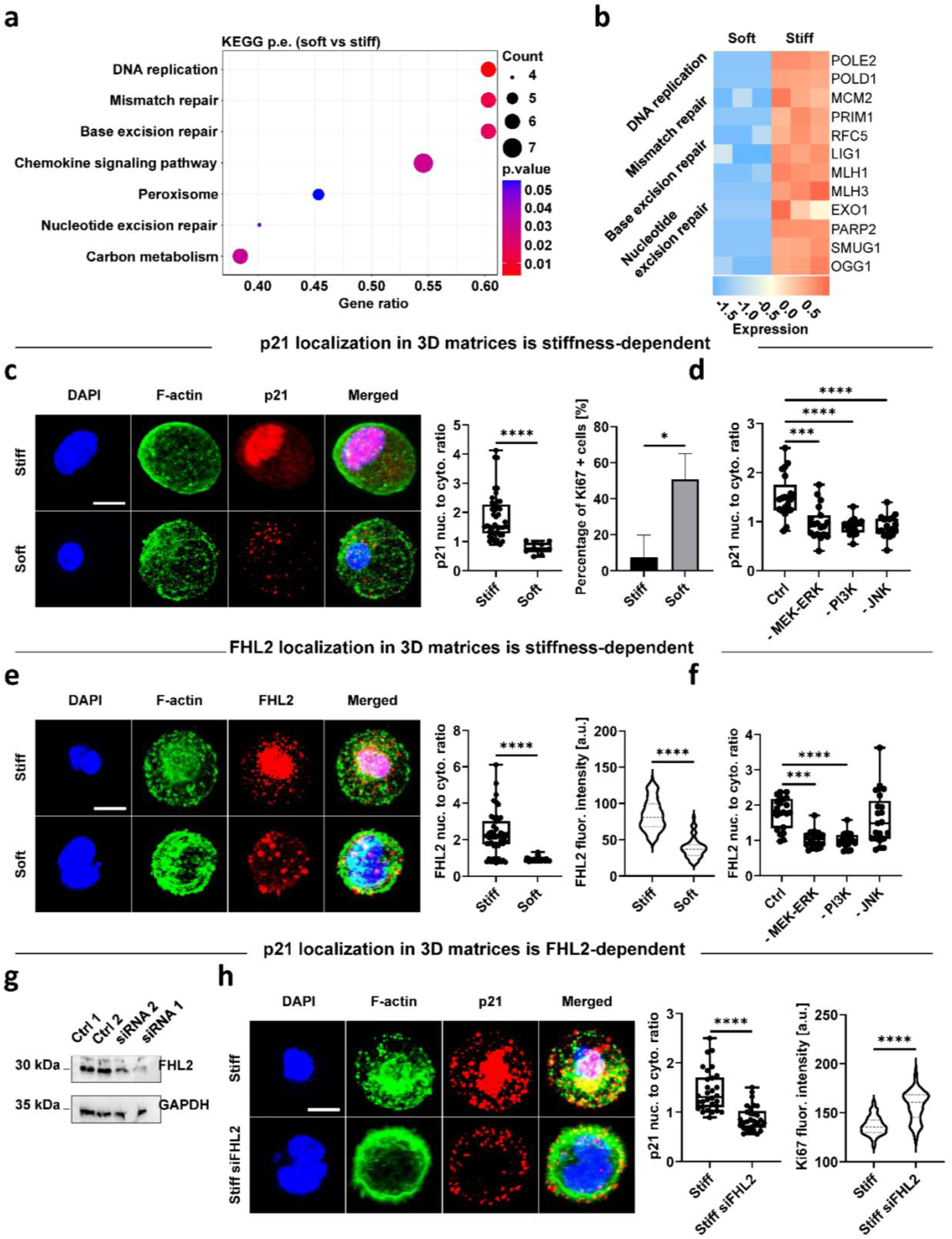
p21 and FHL2 localization in 3D matrices is stiffness-dependent and FHL2 mediates p21 localization and Ki67 expression. **a)** Selected top differentially regulated pathways from GSEA based on differentially expressed genes between cells grown in soft vs. stiff alginate hydrogels. **b)** Heat map of differentially expressed genes (soft vs. stiff) from DNA damage and repair pathways. **c)** Representative confocal images of p21 localization and quantification of its nuclear-to-cytoplasmic ratio (n=3 gels for 10 to 42 cells) for MDA-MB-231 cells within 3D stiff and soft alginate hydrogels after 7 days of encapsulation (blue=DAPI, green=F-Actin, red=p21). Scale bar equals 10 μm, Mann-Whitney *U*-test, ****p<0.0001. Fraction of Ki67-positive cells (n=3 gels for 28 to 628 cells), student’s *t*-test, *p<0.05. **d)** Quantification of p21 localization in 3D alginate stiff hydrogels after 7 days in the absence or presence of indicated inhibitors (n≥19 cells). Student’s *t*-test with respect to control group (3D alginate stiff), ***p<0.001 and ****p<0.0001. **e)** Representative confocal images of FHL2 localization and quantification of its nuclear-to-cytoplasmic ratio (n=3 gels for 36 to 46 cells) for MDA-MB-231 cells within 3D stiff and soft alginate hydrogels after 7 days of encapsulation (blue=DAPI, green=F-Actin, red=FHL2). Scale bar equals 10 μm, Mann-Whitney *U*-test, ****p<0.0001. FHL2 fluorescence intensity (f.i.) (a.u.=arbitrary unit) per cell (n=3 gels for 59 to 151 cells) Mann-Whitney *U*-test, ****p<0.0001. **f)** Quantification of FHL2 localization in 3D alginate stiff hydrogels after 7 days in the absence or presence of indicated inhibitors (n≥18 cells). Student’s *t*-test with respect to control group (3D alginate stiff), ***p<0.001 and ****p<0.0001. **g)** Immunoblot showing FHL2 levels in MDA-MB-231 cells treated with two different anti-FHL2 siRNA oligos compared to untreated (control 1) or scramble (control 2) groups. GAPDH, glyceraldehyde-3-phosphate de-hydrogenase. **h)** Representative confocal images of p21 localization and quantification of its nuclear-to-cytoplasmic ratio (n=3 gels for 30 cells) for MDA-MB-231 normal and siRNA-FHL2 silenced cells within 3D stiff alginate hydrogels (blue=DAPI, green=F-Actin, red=p21). Scale bar equals 10 μm, Mann-Whitney *U*-test, ****p<0.0001. Ki67 f.i. (a.u.=arbitrary unit) per cell (n= 74 to 260 cells), Mann-Whitney *U*-test, ****p<0.0001.

Previous work on 2D cultures had found that, in MDA-MB-231 cells, p21 is induced in response to treatment with DNA damaging drugs, and this is mediated by FHL2 in a p53-independent manner (*46*). Notably, FHL2 was shown to be stably expressed in only a small subset of BCC lines with MDA-MB-231 being among the highest (*46*). Nonetheless, FHL2 localization and functional role in physiologically-relevant 3D cultures had not been addressed. We therefore investigated the nuc./cyto. ratio of FHL2 in 3D stiff compared to soft alginate hydrogels. Surprisingly, and in stark contrast to 2D cultures, where soft 2D substrates lead to nuclear accumulation of FHL2 (*48*), we observed a significantly higher expression and nuc./cyto. ratio of FHL2 in 3D stiff compared to soft alginate hydrogels (Fig.5e). This effect was mitigated by inhibiting MEK/ERK and PI3K/Akt pathways but not JNK (Fig.5f, Supp.Fig.S10). Importantly, knock-down of FHL2 revealed a decrease in p21 nuclear localization in 3D stiff hydrogels (Fig.5g, h), as well as an increase in proliferation activity (Ki67; Fig.5h), pointing to FHL2 being upstream of p21. The correlated expression of p21 (*cdkn1a*) and FHL2 is also found in tumor samples of patients with ER-breast cancer (Supp.Fig.S11).

### FHL2 knockdown sensitizes cells to chemotherapy

We then investigated whether FHL2 knockdown sensitizes cells to chemotherapy. After FHL2 silencing, cells were encapsulated for 3 days, followed by 2-day drug exposure (Fig.6a). FHL2-silenced cells in 3D alginate stiff displayed higher viability and metabolic activity than wild-type MDA-MB-231 when untreated (vehicle) (Fig.6a). Upon exposure to standard 0.01mM and higher doses of paclitaxel in a concentration-dependent fashion (≥0.01mM), a significant reduction in viability and metabolic activity was observed compared to wild type cells (Fig.6a), pointing to a chemo-sensitizing effect upon FHL2 knockdown.

**Fig. 6.**
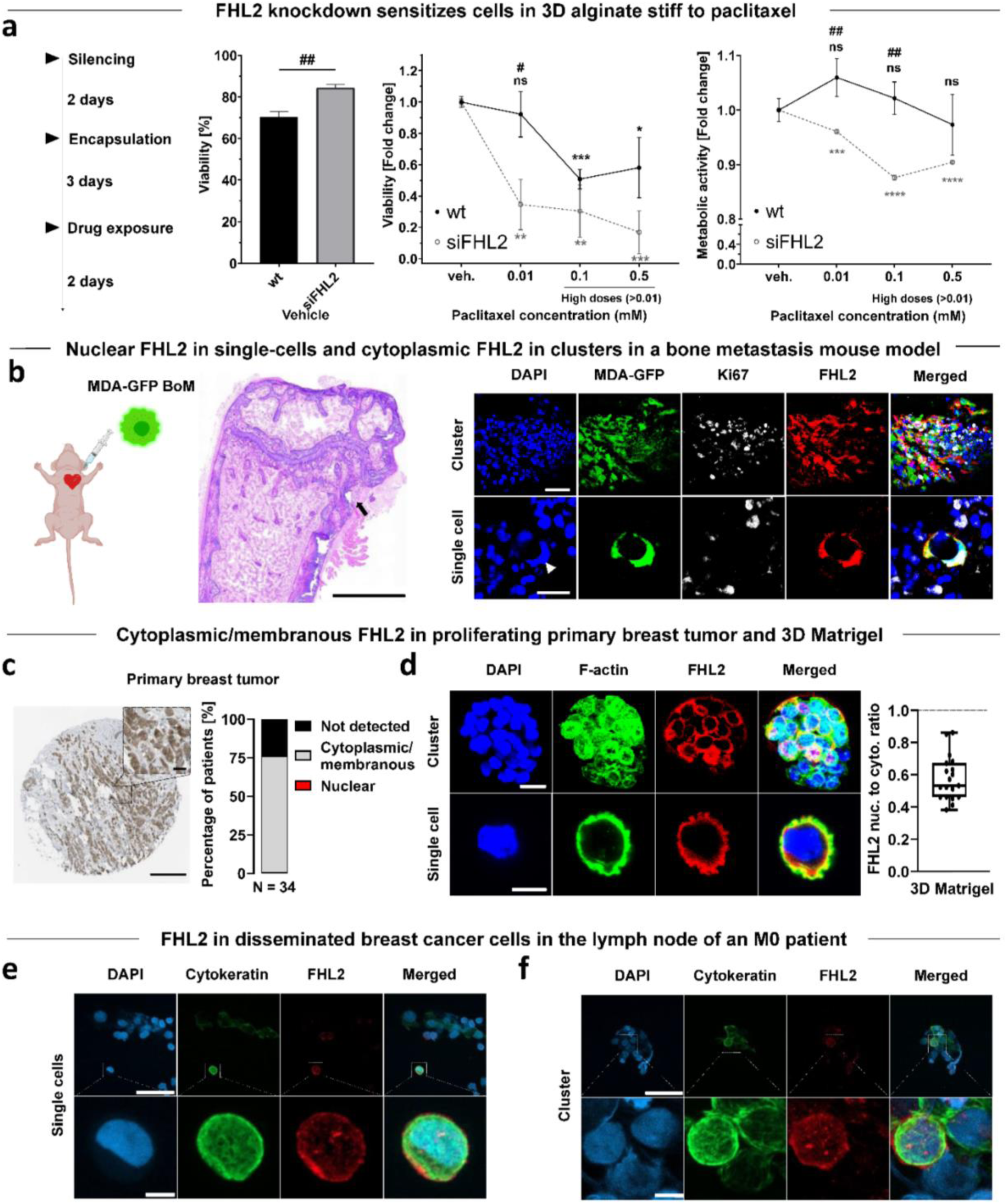
FHL2 knockdown sensitizes cells to chemotherapy. FHL2 expression and localization in murine tissue of a breast cancer bone metastasis model, in human primary breast tumor and in early DCCs of an M0 patient. **a)** FHL2-silenced MDA-MB-231 cells were encapsulated for 3 days and exposed to the standard 0.01mM and higher doses of paclitaxel (0.01 to 0.5 mM) for 2 days, after which viability and metabolic activity (Presto Blue) measurements of normal and siRNA-FHL2 silenced cells within 3D alginate stiff were performed. Percentage bars of viability (calcein+) for the vehicles and fold changes in viability and metabolic activity with respect to the vehicle and between groups were measured. * for statistical analysis with respect to the vehicle of each gel (untreated groups) and # for statistics between groups. Student’s *t*-test (n=3 hydrogels), */#p≤0.05, **/##p≤0.01, ***/###p≤0.001. **b)** Intracardiac injection of GFP-tagged MDA-MB-231-1833 BoM cells in mice with representative Hematoxylin and Eosin (H&E) staining of the femur showing an osteolytic lesion (arrowhead). Representative confocal images of Ki67 (proliferation) and FHL2 localization of GFP-tagged MDA-MB-231-1833 BoM cells as clusters and single cells. Scale bars equal 1000 μm, 25 μm and 20 μm for H&E overview, cluster and single cell images, respectively. **c)** Representative immunohistochemical image of FHL2 localization in a female patient (50 years old) with breast ductal carcinoma. Fraction of FHL2 localization in 34 patients with primary breast ductal carcinoma. Data obtained from The Human Protein Atlas^90^. Scale bar equals 20 µm. **d)** Representative confocal images of FHL2 localization in spheroids and single MDA-MB-231 cells, both kinds encapsulated within the same 3D Matrigel after 7 days of encapsulation and combined quantification of its nuclear-to-cytoplasmic ratio (n=19 cells) (blue=DAPI, green=F-Actin, red=FHL2). Scale bar equals 20 μm for spheroid image and 10 μm for single cell image. **e, f)** Confocal images of cytokeratin (epithelial marker) and FHL2 (in single cells and within clusters) of human breast cancer cells disseminated in the sentinel lymph node of an M0 patient. Scale bars equal 50 μm and 5 μm for the wide and zoomed images, respectively.

### FHL2 expression and localization in murine tissue, human primary breast tumor and human early DCCs

Finally, to assess the implications for human disease, we investigated FHL2 expression and localization in a preclinical mouse model with metastatic breast cancer and in human primary tumor biopsies. Remarkably, single MDA-MB-231 cells (bone-tropic 1833 subclone) spontaneously disseminated to the mouse femur revealed nuclear FHL2, as opposed to proliferative clusters with diffuse cytoplasmic FHL2 (Fig.6b, Supp.Fig.12). Similarly, human tissue biopsies of breast tumor reveal a primarily cytoplasmic/membranous localization of FHL2 (Fig.6c), matching our observation in 3D Matrigel (Fig.6d). Furthermore, we investigated human early DCCs of an M0 breast cancer patient, with only microscopy evidence of metastasis and no clinical or radiographic evidence of distant metastasis. DCCs (cytokeratin+, an epithelial marker) in the sentinel lymph node displayed varying FHL2 signal intensity suggestive of functional heterogeneity (Fig.6e, f).

To sum up, ligand-rich, proliferation-permissive microenvironments maintain FHL2 in the cell membranous regions, whereas in low or non-adherent microenvironments the increase in mechanical confinement (stiffness) results in FHL2 translocation to the nucleus, leading to high p21 nuclear expression, cell cycle arrest and chemo-resistance (Fig.7).

**Fig. 7.**
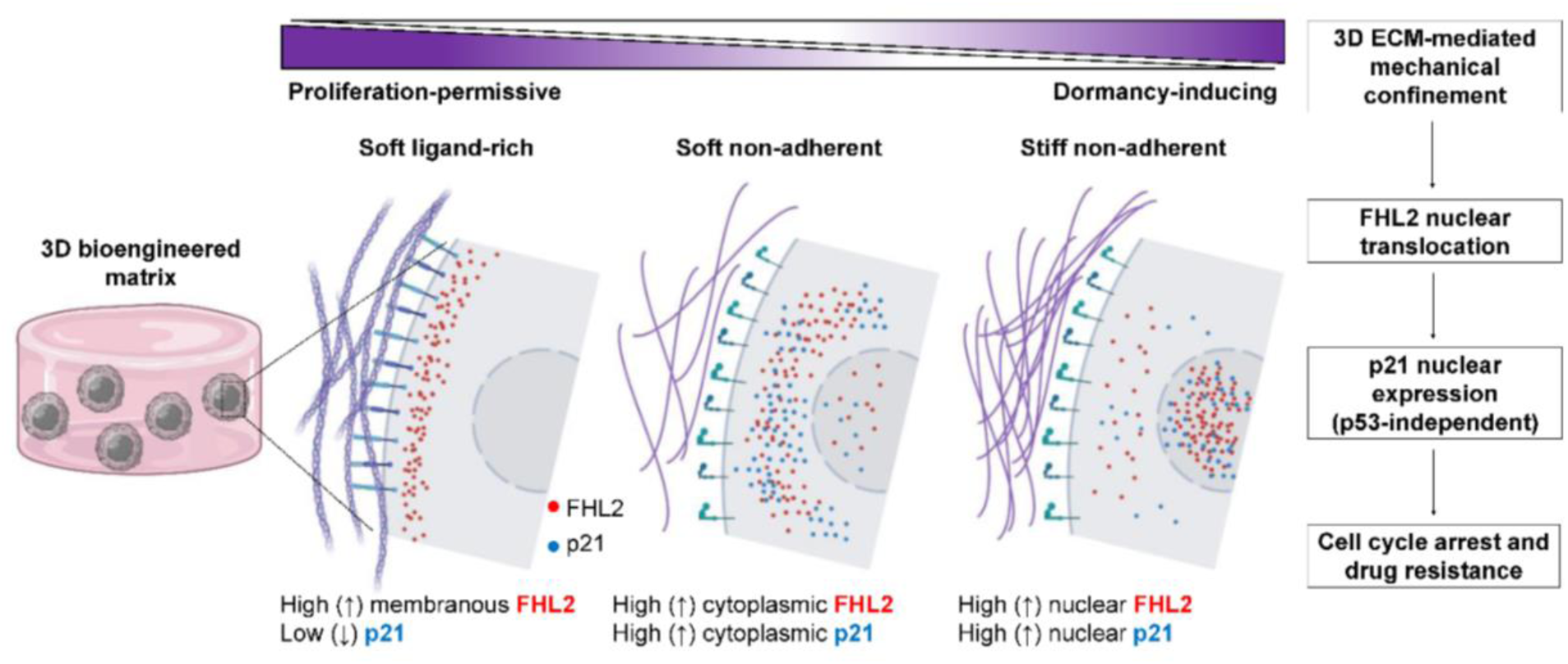
Proposed stiffness-mediated FHL2 signaling mechanism in 3D bioengineered matrices inducing cancer cell dormancy and drug resistance. In proliferation-permissive, ligand-rich microenvironments such as basement membranes, stronger adhesion sites maintain FHL2 in the membranous regions. In low or non-adherent microenvironments, the increase in mechanical confinement (stiffness) results in FHL2 translocation to the nucleus, leading to high p21 nuclear expression and cell cycle arrest (graphical illustration created with BioRender.com).

## Discussion

In this study, we use a 3D dormancy-inducing engineered matrix to investigate a mechanism of chemoresistance in mechanically-induced dormant BCCs. Systematic tuning of three variables (adhesion via RGD, degradability through MMP-sensitive crosslinkers and stiffness) enables generation of different fractions of cell populations with defined and controllable cell cycle status. Here, we harness this approach and reveal a stiffness-dependent low P-ERK^low^:P-p38^high^ dormancy phenotype, along with Ki67-expression. We further validated an upregulation of CDK inhibitors p21 and p27, together with a nuclear localization of the FHL2 protein which leads to p53-independent nuclear expression of p21 as a dormant and resistance-conferring mechanism (Fig.7).

We have demonstrated that the nuclear localization of p21 in 3D matrices is FHL2 dependent. We show this indirectly by (i) knocking down FHL2 expression using siRNA, (ii) quantifying the p21 nuclear to cytoplasmic localization and showing a shift towards cytoplasmic localization (Fig.5h), accompanied by (iii) an increase in Ki67 proliferation marker (Fig.5h). Furthermore, we show that knocking down FHL2 has a chemo-sensitizing effect to paclitaxel (Fig.6a). In our previous study, we could show with Brillouin microscopy that the nucleus and cytoplasm of cells in stiffer matrices become denser (*49*). One possibility is that stiffer matrices induce densification of nucleus and cytoplasm and FHL2 could move directly to the nucleus due to passive diffusion, as FHL2 is considered small enough to diffuse through nuclear pores (*48*). However, a deeper molecular analysis of FHL2 and p21 protein shuttling to the nucleus in 3D matrices was beyond the scope of this study.

We further show that the expression and localization of both p21 and FHL2 in 3D is strikingly different from previously-reported 2D studies, where FHL2 shuttling to the nucleus was shown to be substrate rigidity-dependent, such that on 2D soft surfaces (as opposed to stiff substrates where mechanical tension is higher) it would translocate to the nucleus and induce p21 expression (*48*). In contrast, in 3D ligand-rich environments such as in basement membrane matrices or primary breast tumor sites, FHL2 accumulates in adhesion sites (*48*), whereas loss of adhesion (occurring in intrinsically non-adherent materials such as alginate), results in FHL2 release from sub-membranous regions and shuttling to cytoplasmic (soft alginate) or nuclear (stiff alginate) regions. This dimensionality-dependent behavior across adhesion sites has been similarly reported for YAP mechanotransduction in the context of breast tumor progression (*50*) and stem cell fate (*51*). Thus, it is critical to take into account the context-dependent role of the FHL2-p21 signaling axis, given its potential dual oncogenic vs. suppressive function (*46*, *52*).

FHL2 has been previously shown to concentrate at RNA polymerase (Pol) II sites (*48*), inducing c-Jun and IL8 transcription. This is reflected in our sequencing data by the strong enrichment of RNA Pol II associated genes, with JUN and IL8 (CXCL8) being among the top ten (out of 9427) differentially regulated genes in the alginate stiff group (Fig.4b-e).

Omission of adhesion ligands in our dormancy-inducing matrix was intentional, stemming from recent *in vivo* observations (*11*) where single DCCs were shown to reside in a dormant state with expressed but not engaged integrin receptors. Other studies have similarly observed DCCs expressing low levels of adhesion molecules (*24*), and integrin activation being a central mediator leading to their awakening (*53*–*56*). We enhance the idea that the survival of our non-proliferative BCCs in a stiff non-degradable and non-adherent matrix is mainly regulated by the p-ERK^low^:p-38^high^ dormancy ratio and the consequent p21 and p27 upregulation (Supp.Fig.S6), in agreement with previous *in vivo* observations (*3*, *38*, *57*). As stated, this cascade leads to p21 nuclear localization, cell cycle arrest and drug resistance (Fig.7). BCCs in G_0_/G_1_ phase revealed a resistance to low doses of cell cycle-specific paclitaxel (Fig.3). Interestingly, silencing FHL2 had a chemo-sensitizing effect to paclitaxel, with a significant decrease in both viability and metabolic activity even at higher doses of paclitaxel (Fig.6a).

All these effects were only characterized for the commonly-used metastatic BCCs, which have previously shown short-lived growth arrest and recovery capacity upon serum starvation (*31*). However, prior work has established suitable cell lines that can enter growth arrest and recover after 8 weeks of serum starvation, such as D2.0R, a murine triple negative dormant breast cancer cell line considered highly serum-dependent (*31*, *33*). In this work, we demonstrate the potential of our biomaterial-based model to drive human metastatic cell lines into dormancy within a few days, being able to recapitulate P-ERK^low^:P-p38^high^ dormancy phenotype along with Ki67-/p21+/p27+ expression. Although this does not fully simulate long-term growth arrest and recovery phenotypes (*28*), this platform could be advantageous to induce patient-derived cells into a dormant phenotype within a few days, mainly dominated by matrix stiffness and regardless of their origin and heterogeneous behavior.

To further support our findings, we investigated BCCs disseminated to the bone in a metastasis mouse model, human tissue biopsies of primary breast cancer and early DCCs of an M0 breast cancer patient. Here we noted nuclear FHL2 localization in single dormant DCCs (Fig.6b) as opposed to cytoplasmic FHL2 corresponding to high proliferation activity (Ki67+) in a mouse model of breast cancer metastasis (Supp.Fig.12). Likewise, cytoplasmic/membranous FHL2 is observed in human primary breast tumor biopsies (Fig.6c) matching the findings of proliferating cancer cells in 3D Matrigel (Fig.6d). This is in agreement with screenings of normal and malignant human breast tissues revealing higher FHL2 expression for the latter, as shown in previous studies (*58*, *59*) and UALCAN database, while high FHL2 expression in ER- patients is associated with lower relapse-free survival rate (Supp.Fig.13). A number of disseminated BCCs (cytokeratin+) were found in the sentinel lymph node both as single cells (Fig.6e) and within cell clusters (Fig.6f). We noted a substantial heterogeneity among the DCCs, further supporting the notion that various phenotypes existing in patients might require different targeting approaches.

From a translational perspective, the complexity associated with detection, isolation and analysis of DCCs in asymptomatic patients poses technical challenges for drug and target discovery in cancer dormancy. Consequently, our bioengineered platform offers new opportunities for drug screening of single dormant DCCs in a short period of time. In particular, our biomaterial-based approach can induce a state of growth arrest of single BCC upon 5 days encapsulation in 3D alginate stiff hydrogels. This is an effective approach to generate a large pool of multiple individual growth-arrested BCCs within a few days. Scalability of high throughput drug screening could be implemented by using multiwell plates to introduce gels and test different markers or treatments as shown previously (*33*, *35*). Others have used alternatives such as 3D printing of large batches of hydrogels (*60*), or robotic arms to automatically inject small 3D microgels into multiwell plates, overcoming the limits of 2D high-throughput work flow techniques (*61*). The mechanically-induced single dormant DCCs could be characterized by multiomics analyses for target discovery and identification of biomarkers related to dormancy and survival. The identified targets and pathways could be validated using patient-derived cells (*60*). High-throughput chemical biology and functional genomics approaches (e.g. Clustered Regularly Interspaced Short Palindromic Repeats (CRISPR) screening) could be combined with 3D phenotypic high-content screening as well as the emerging omics profiling technologies at scale to identify/validate targets and to screen large compound libraries. This could facilitate the discovery of novel lead structures with the potential to selectively eradicate dormant DCCs and/or prevent their reactivation.

To conclude, we identify a stiffness-mediated FHL2 signaling mechanism in 3D bioengineered matrices inducing BCC dormancy and drug resistance (Fig.7), which recapitulates several aspects of patient-derived DCCs and mouse models of metastasis. Further, this approach can be exploited to generate large populations of single growth-arrested cells for simple, fast and potentially scalable drug screening, since these are known to be rare and inaccessible in large numbers from clinical settings.

## Materials and Methods

### Synthesis of peptide-crosslinked alginate hydrogels

Norbornene-modified alginate hydrogels with thiol-ene crosslinking were synthesized as previously described (*39*). Briefly, high molecular weight (>200 kDa; HMW), high guluronic acid, sodium alginate (Pronova MVG; NovaMatrix) was dissolved at 1% w/v in 0.1 M 2-(N-morpholino)ethanesulfonic acid (MES; Sigma-Aldrich), 0.3M NaCl (EMD Millipore) buffer (pH 6.5) overnight at room temperature. N-hydroxysuccinimide (NHS; Sigma-Aldrich), followed by 1-ethyl-3-(3-dimethylaminopropyl)-carbodiimide hydrochloride (EDC; Sigma-Aldrich) were added drop-wise at 5000 molar equivalents to the alginate solution while stirring. To functionalize the alginate backbone with norbornene functional groups, 5-norbornene-2-methylamine (TCI Deutschland GmbH) was added at a theoretical degree of substitution (DStheo) of >300 molecules per alginate chain. The final concentration of both reactions was 0.6% w/v and was run while stirring at 700 rpm at room temperature for 20 h. Next, the solution was quenched by adding hydroxylamine hydrochloride (Sigma-Aldrich), followed by dialysis (Spectra/Por 6, MWCO 3.5 kDa; Spectrum) against a salt gradient (6 g/L to 0 g/L; Sigma-Aldrich) in ddH_2_O for 3 days with 3-4 changes per day. The solution was then purified with activated charcoal (Sigma-Aldrich), sterile-filtered (0.22 μm; Steriflip-GP; Merck) and lyophilized. To assess the actual degree of substitution, NMR measurements were performed using an Agilent 600 MHz PremiumCOMPACT equipped with Agilent OneNMR Probe (256 scans), with samples dissolved at a final concentration of 1.5% w/v in deuterium oxide (D_2_O) (Supp.Fig.S1b, c).

### Casting of peptide-crosslinked alginate hydrogels

For hydrogel casting, the VPMS↓MRGG sequence was chosen as the enzymatically-degradable peptide crosslinker (↓ denotes the MMP cleavage site), ordered from WatsonBio Sciences at 98% purity with trifluoroacetic acid removal (Supp.Fig.S1d). Double-cysteine containing dithiothreitol (DTT, Sigma-Aldrich, 43816) was used as the non-degradable crosslinker. Before casting, norbornene-modified alginate and the photoinitiator (Irgacure 2959; Sigma-Aldrich) were dissolved in phosphate-buffered saline (PBS) overnight at 50 °C under shaking. Alginate and photoinitiator concentrations were kept at 2 and 0.5% w/v, respectively. The concentration of the crosslinker was changed to yield hydrogels with different mechanical properties. After mixing, the solution was pipetted with positive-displacement pipettes on a glass plate and covered with a dichloromethylsilane-coated glass slide (≥99.5%; Sigma-Aldrich). To initiate thiol-ene crosslinking, the mixture was exposed to UV light (365 nm at 10 mW/cm^2^, OmniCure S2000) by placing the gel sheet in a custom-built curing chamber for at least 3min. Gels were then punched out using biopsy punches of 4-6 mm (Integra Miltex) and washed with PBS or medium until further use. To render norbornene-modified alginate adherent, a thiol-containing RGD sequence (CGGGGRGDSP; Peptide 2.0) was added to the gel precursor mix (at 0.95 mM) to bind residual norbornene groups via UV-mediated thiol-ene chemistry. For 3D cell encapsulation, a cell suspension (5×10^5^ cells/ mL) was added to the precursor solution and mixed before UV exposure. Matrigel (356237, Corning) was used at 100% w/v concentration (∼ 10mg/mL).

### Mechanical characterization

Rheology measurements were performed to calculate the elastic modulus (E) of pre-formed gels. Norbornene-modified alginate (2% w/v) hydrogels with different stiffness were cast by changing the DTT crosslinker concentration (0.05-1 mg/mL). After gel equilibration overnight in PBS, frequency sweep measurements were performed from 0.01 to 10 Hz at 1% shear strain with an 8 mm parallel plate (PP08, Anton Paar) using a rheometer (Physica MCR 301; Anton Paar), while keeping the temperature at 25 °C with a Peltier cooling module. The initial E modulus was then calculated as follows: *E* = 2*G*(1 + *ϑ*) and 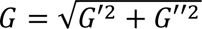 where *G*, *G*′ and *G*′′ are the shear, storage and loss moduli, respectively, and *ϑ* the Poisson’s ratio with a value of 0.5 for hydrogels (*62*).

### Cell culture

MDA-MB-231 (HTB-26; ATCC) and MDA-MB-231-1833 BoM (provided by Dr. Joan Massagué and purchased from the Antibody and Bioresource Core Facility of the Memorial Sloan Kettering Cancer Center, USA) cell lines were cultured as models for triple-negative breast cancer and MCF-7 (HTB-22; ATCC) human breast cancer cell line for the luminal subtype. The subclone 1833 is a bone tropic human cell line deriving from a metastasis formed by MDA-MB-231 cells hosted in a mouse (*63*, *64*), which in turn are MDA-MB-231 human epithelial breast cancer cells stably transduced with a lentivirus expressing a triple-fusion reporter (*65*). These three cell lines were cultured in low glucose Dulbecco’s Modified Eagle’s Medium (D6046; Sigma-Aldrich) with 10% v/v fetal bovine serum (S0615, Sigma-Aldrich) and 1% penicillin/streptomycin (Gibco). For MCF-7 cells, 0.1% insulin (I1882, Sigma-Aldrich) was added. Cells were then incubated in a 5% CO_2_ environment at 37 °C and passaged every 3-5 days.

### Viability and metabolic activity characterization

Live and dead cells were stained with 1.6-4 mM calcein AM (C125400; TRC) and 4 mM ethidium homodimer-1 (EthD-1 L3224; ThermoFisher), respectively. To measure metabolic activity, Presto blue (ThermoFisher) was used. Briefly, after 1-7 days of incubation, cell culture medium was replaced with 10% Presto blue reagent in DMEM and the plate incubated for at least 4 h at 37 °C. Next, 100 µL of the supernatant was transferred to a black 96 well plate (655892, Greiner Bio-one) and fluorescence emission was measured at 590 nm (Cytation5, BioTek) and at 580-640nm (Glomax, Promega).

### Lentiviral particle production, transduction and generation of FUCCI2 breast cancer cell lines

As previously reported (*66*), FUCCI2 vectors mCherry hCdt1(30/120)/pCSII EF MCS (DDBJ/EMBL/GenBank, AB512478) and mVenus hGeminin(1/100)/pCSII EF MCS (DDBJ/EMBL/GenBank, AB512479) were purchased from the Riken Brain Science Institute (Japan) and used to generate lentivirus particles by co-transfecting HEK 293TN cells (System Biosciences) with packaging (psPAX2, Addgene plasmid, #12260) and envelope (pMD2.G (Addgene plasmid, #12259) plasmids. The supernatant was collected by centrifuging (Beckman L7-55 with SW32Ti rotor) at 22,000 rpm for 3h at 4 °C. MDA-MB-231 and MCF-7 breast cancer cell lines were first transfected with mVenus hGeminin (1/110) (multiplicity of infection (MOI) of six and five, respectively) followed by mCherry hCdt1 (30/120) at an MOI of three. Verification of successful transduction and subsequent sorting of stably expressing mVenus and mCherry cell lines were performed using flow cytometry (FACSAriaTM II, Becton Dickinson).

### Inhibition, RNA interference and drug response experiments

For inhibition experiments, the following small molecule and inhibitors were used: MEK/ERK inhibitor PD98059 (10 mM, Focus Biomolecules), JNK inhibitor SP600125 (10 mM, Focus Biomolecules) and PI3K/Akt inhibitor LY294002 (10 mM, Focus Biomolecules). The inhibitors were mixed with culture medium and refreshed every 2-3 days. To knock down FHL2, MDA-MB-231 cells were transfected with two different FHL2-targeting small interfering RNAs (Silencer siRNA, Invitrogen) or non-targeting siRNA (Stealth RNAi negative control, Invitrogen) using Lipofectamine RNAiMAX Reagent (Invitrogen), diluted in Opti-MEM reduced serum medium (Gibco) for a final concentration of 10-20 nM. Cells were transfected 2-3 days before encapsulation, following the manufacturer’s instructions.

To assess MDA-MB-231 response to chemotherapy, Paclitaxel (Tokyo Chemical Industry), Gemcitabine (#710786.6, Aurouitas), 5-Fluorouracil (#603544.3, Accord) and Carboplatin (#670568.2, Accord), commercial brands kindly provided by the Department of Pharmacy in Onkologikoa (San Sebastian, Spain), were diluted at 0.001, 0.005, 0.01, 0.1 and 0.5 mM. After 3 days of encapsulation, dilutions were added to the culture medium of the gels, targeting encapsulated cells with 2-day drug treatment. Experiments performed with paclitaxel at the highest concentration contained around 1% (v/v) of DMSO, which was used as vehicle and didn’t show any negative effect on cell viability. Excipients and dissolvent of drugs are presented in Supplementary Table S1.

### Immunofluorescence staining

Gels with encapsulated cells were washed (max. 3x) with PBS and fixed with 4% paraformaldehyde (PFA, Boster, #AR1068) for 30 min at room temperature, (3x) washed with 3% w/v bovine serum albumin (BSA, Sigma Aldrich), permeabilized with 0.1 % w/v Triton X 100 (Sigma-Aldrich, #T8787) for 15 min with a final washing (3x) step with 3% BSA. Whole-mount primary antibody staining was performed at 4 °C overnight, diluted in a 3% BSA+0.1% Triton X 100 buffer solution under mild shaking. The gels were then washed (3x) with 3% BSA and incubated with secondary antibody at 4 °C overnight with the same dilution buffer under mild shaking. The samples were finally imaged after a last washing step with PBS (3x). The primary antibodies and respective concentrations used in this study are the following: anti-Ki67 (2 µg/mL, ab15580, abcam), anti-p21^Cip1/Waf1^ (0.48 µg/mL, 2947S, Cell Signaling), anti-FHL2 (0.5 µg/mL, HPA006028, Sigma/Prestige Antibodies), anti-phospho-p38 (1:200, #9216S, Cell Signaling) and anti-phospho-p44/42 MAPK (p-ERK1/2) (1:200, #9101S, Cell Signaling). The secondary antibodies are: Alexa Fluor 405 (10 µg/mL, ab175651, abcam), Alexa Fluor 488 (1:500, #44128878S, Cell Signaling) and Alexa Fluor 647 (4 µg/mL, A21244, Invitrogen and 1:500, #4410, Cell Signaling). To stain for nucleus and cytoskeleton, DAPI (1:1000, Roche) and Alexa Fluor 488 Phalloidin (1:50, Invitrogen) were used.

### Immunoblot

Cells were first lysed with RIPA buffer (Abcam) for 5 min on ice and then collected in Eppendorf tubes. Protein concentration was calculated following Bradford assay (Bradford reagent, Sigma-Aldrich). A 1:50 mixture of β-mercaptoethanol (Acros Organics) and 2x Tricine sample buffer (Bio-Rad) was added at a 1:1 volume to the cell lysates. The mixture was then heated to 95 °C for 5 min and loaded into pre-cast polyacrylamide gels (Mini-PROTEAN TGX, Bio-Rad) for electrophoresis in TGS buffer (Bio-Rad) at 100 V for 30 min, then 200 V for 30 min (Mini-PROTEAN Tetra System, Bio-Rad). Proteins in the gels were then blotted onto to a PVDF membrane (BioTrace, PALL) at 150 V for 90 min in ice-cold transfer buffer (1:5 volume of 100% Methanol in TGS buffer). Next, the membrane was transferred to TBST buffer (150 mM NacL, 20 mM Tris-base, 0.1% Tween20), blocked in 5% BSA in TBST buffer for 1 h under gentle shaking, washed in TBST (3x) and incubated with primary antibodies diluted in 1% BSA in TBST overnight at 4 °C. After washing with TBST (3x), the membrane was incubated with an HRP-linked secondary antibody for 1 h at room temperature, after which the chemoluminescence substrate (SuperSignal West Pico PLUS, Thermo Scientific) was added before visualizing it with a blot imager (G:BOX Chemi XX6/XX9, Syngene). GAPDH (1:1000, 2118, Cell Signaling) was used as housekeeping gene and HRP-linked antibody 7074 (1:1000, Cell Signaling) as secondary antibody.

### Cell retrieval from 3D gels

MDA-MB-231 cells were retrieved from 3D alginate stiff, soft and 3D Matrigel. Briefly, alginate hydrogels were incubated with alginate lyase (2mg/mL, Sigma Aldrich, A1603) in HBSS (ThermoFisher, 14185045) for 30 min at 37 °C with three mixture steps every 10 min. For Matrigel, pre-cooled (4°C) HBSS was added and mixed for 5-10 min at room temperature, followed by incubation with Trypsin/EDTA (0.05%/0.02%, PAN Biotech, P10-023100) for 5 min at 37 °C. Cells were then washed twice with HBSS for 5 min. Next, retrieved cells were stained using a LIVE/DEAD fixable cell staining method (ThermoFisher, L34969) following the manufacturer’s instructions, and viable cells were sorted via FACSAria II flow cytometer (Becton Dickinson). Approximately 1500 sorted cells per sample were then lysed using pre-cooled (4 °C) RLN buffer (0.05 M Tri-HCl pH 8.0, 0.14 M NaCl, 0.0015 M MgCl_2_, 0.5 % v/v Nonidet P-40/IGEPAL (Sigma Aldrich, 56741) and 1000 U/mL RNase inhibitor (Promega, N2615), 0.001 M DTT) on ice for 5 min. The lysates were then centrifuged with a pre-cooled centrifuge for 2 min at 300 x g, the supernatants transferred to -80 °C and stored until further processing.

### RNA sequencing and analysis

Samples were resuspended in 5 µl 1X NEBNext Cell Lysis Buffer of the NEBNext Single Cell/Low Input RNA Library Prep Kit (NEB, E6420). After 5 min incubation at RT, 2.5X RNA XP Clean Up Beads (Beckman Coulter, A63987) were added to the sample by pipetting up and down. Then, samples were incubated for 5 min at RT, the beads were pelleted on a magnet, the supernatant was removed, the samples were washed twice with 80% ethanol while remaining on the magnet, and finally the beads were resuspended in 8 µl nuclease-free water. The reverse transcription was conducted according to the manual of the NEBNext Single Cell/Low Input RNA Library Prep Kit in the presence of the beads. Next, the beads were pelleted on the magnet and the supernatant was used for the following PCR amplification as described in the manual. Twenty-one PCR cycles were applied, because the RNA concentration was below the detection limit of the Qubit RNA HS Assay Kit (Thermo Fisher, Q23852). The cleanup, quality control, fragmentation, and adapter ligation steps were performed as described in the kit manual. NEBNext Multiplex Oligos for Illumina (Index Primer Set 1) (NEB, 7335S) were used for the final PCR amplification. The fragment length distribution of the final libraries was determined using a 2100 Bioanalyzer Instrument (Agilent) with a High Sensitivity DNA kit (Agilent, 5067-4626). The libraries were quantified by qPCR with the KAPA Library Quantification Kit (Roche, KK4854). The nine samples were divided into two pools (one pool of four samples, the other of five samples) with equimolar amount. The pools were sequenced on an Illumina MiSeq with 2 x 150 bp.

After demultiplexing, raw FASTQ data were given to an in-house mRNA analysis pipeline 0.9.5.5. BBDuk 38.76 (*67*) was used to trim the raw sequence data from 3D Matrigel, 3D alginate soft and stiff, deleting any residual adapter sequences and low-quality bases at the ends of each read. BioBloom Tools 2.0.13 (*68*) was used to decontaminate reads from the genomes Mus musculus (mm38), Escherichia coli (BL21), Mycoplasma pneumoniae (M129), Sphingobium sp. (SYK-6), Bradyrhizobium japonicum (USDA 110), Pichia pastoris (GS115), Malessia globosa (CBS 7966), Aspergillus fumigatus (Af293), and a set of viral genomes (RefSeq, 5k+ genomes). All reads that did not map exclusively to the transcriptome of hg38 (GENCODE version 27, GRCh38.p10) were labeled as potentially contaminated and were removed from further processing. FastQC 0.11.9 (*69*) was used to evaluate sequence quality per sample before as well as after trimming and decontamination. In addition, all samples were examined as a collective with MultiQC 1.8 (*70*). Following that, the cleaned sample reads were aligned to the hg38 reference genome using STAR 2.5.1b (*71*). Using featureCounts from Subread 2.0.0, uniquely mapping reads were counted per gene and sample (*72*). Further quality criteria were evaluated, including library complexity (using Preseq 2.0.3 (*73*)) and the genomic origin of the reads and the 5’-3’-bias (both using QualiMap 2.2.2d (*74*)). The final counts table of 9 samples were utilized for differential expression analysis.

All the subsequent steps after the count tables were performed in R programming language 4.0.2 (2020-06-22) (*75*). Matrix table containing gene counts were visualized using Principle Component Analysis (PCA) and t-distributed stochastic neighbor embedding (t-SNE) clustering techniques. For PCA, raw counts were scaled and prcomp function from stats package was employed. tSNE plots were constructed using Rtsne 0.15 (*76*). Subsequent steps including Quality Control (QC) analysis, filtering, normalization, feature selection, scaling, regression of unwanted variables were performed using Seurat package 4.0.2 (*77*). Default parameters were taken for all used functions unless otherwise mentioned. Normalization was done using ‘log.normalize’ method with a scale factor of 1×10^-6^ to obtain logCPM count values and 2,000 most variable genes were used for feature selection. Only mitochondrial genes were regressed out during scaling, cell cycle associated genes did not have an effect on the cell-to-cell clustering. “DESeq2” method was applied to do differential expression analysis between the three groups, soft, stiff and Matrigel and in addition testing of genes was limited to a logFoldChange (logFC) cutoff of 1. Further, the differentially expressed genes were manually filtered for an adjusted *p-value* of 0.05. Finally, volcano plots were constructed using EnhancedVolcano 1.8.0 (*78*). In addition to applying single cell method, Seurat, we also applied pure bulk RNA seq method, DESeq2 (*79*), to perform the data analysis of the single cell pool samples. Results from both methods were highly similar including the list of differentially expressed genes, their respective fold changes and adjusted *p-*value. We chose to report results from Seurat due to the zero-inflated count distribution of our data, which is more typical for single cells than bulk RNA data (Supp.Fig.S4).

Gene Set Enrichment Analysis (GSEA) of significantly Differentially Expressed Genes (DEGs) (adjusted *p*-value < 0.05 and |logFC|> 1) into functional categories were performed using Bioconductor packages 1.30.15. GOseq 1.42.0 (*80*) and clusterProfiler 3.18.1 (*81*) were used, respectively, to conduct Gene Ontology (GO) (*82*) and Kyoto Encyclopedia of Genes and Genomes (KEGG) (*83*) enrichment analyses. All the enrichment plots including GSEA, dot plot and heatmap were made using built-in functions in Seurat package or ggplot2 3.3.3 (*84*). Network maps were processed based on the selected enriched GO categories using Cytoscape 3.8.2 and enrichment Map plugin (*85*).

### RT-qPCR

After RNA quantification using an EPOCH spectrophotometer, 1 µg of RNA was converted to cDNA using the Maxima First-Strand cDNA synthesis kit with incorporated DNAse treatment (#15273796, Thermo Fisher Scientific) according to the manufacturer’s instructions. qPCR was performed using TB Green® Premix Ex Taq™ (Tli RNase H Plus) (#RR42WR, Takara Bio) with a final forward and reverse primer concentrations of 1 µM and cDNA concentrations of 2.5 ng/µl. qPCR was conducted for up to 40 cycles on a CFX 384 PCR system (Bio-Rad). Gene expression levels were determined using the ΔΔCt method and normalized against two stable and common housekeeping genes, ACTB and 18S (*86*). Primers were purchased from Condalab and primer sequences used for qPCR are summarized in Supplementary Table S2.

### Animal experiment

12-week-old female BALB/c nude mice (CAnN.Cg-Foxn1nu/Crl, Charles River, Sulzfeld, Germany) were acclimatized in the animal facility of the Charité-Universitätsmedizin Berlin and housed with ad libitum access to food and water. Mice were injected into the left ventricle of the heart with MDA-MB-231-1833 BoM cells (5×10^5^ cells in 100 µL ice cold PBS), using a 27G needle, under ultrasound guidance (Vevo2100, FUJIFILM VisualSonics Inc., Canada), as previously described (*13*). The animals received Carprosol (CP-Pharma Handelsgesellschaft mbH, Burgdorf, Germany) and Bupresol (CP-Pharma Handelsgesellschaft mbH, Burgdorf, Germany) as analgesic drugs during and after the injection. The animals were anesthetized using isoflurane (CP-Pharma Handelsgesellschaft mbH, Burgdorf, Germany) and the eyes were protected from drying with Pan-Ophtal gel (Dr. Winzer Pharma GmbH, Berlin, Germany). The mouse was sacrificed after 2 weeks by cervical dislocation. The hind limbs were harvested and fixed with 4% paraformaldehyde (PFA) in phosphate-buffered saline (PBS) for 12 h at 4 °C and stored in PBS until further processing. All animal experiments were carried out according to the policies and procedures approved by local legal research animal welfare representatives (LAGeSo Berlin, G0224/18).

For each mouse, bones of one limb were cold embedded at 4°C in poly(methyl methacrylate) (PMMA) (Technovit 9100, Kulzer, Germany), following the manufacturer’s instructions. Briefly, samples were dehydrated in an ascending ethanol series, followed by a xylene washing step, infiltration and embedding in PMMA. The staining was performed on 6 µm thick longitudinal sections using a standard protocol of Haematoxylin and Eosin staining (H & E Rapid kit, Clin-Tech, Surrey, UK). Samples were washed in tap water, followed by staining in Carazzi’s double-strength Haematoxylin. After another wash in tap water, the slides were stained in Eosin, rinsed in tap water, dehydrated and mounted. The stained sections were imaged with a Keyence Digital Microscope (VKX-5550E, Keyence, Germany). For immunostaining of the slides, after deplastification and rehydration, antigen retrieval was performed (sodium citrate pH 6.0/ 0.05% Tween20) at 105°C for 15 min, followed by 10 min quenching of endogenous peroxidase activity with 3% hydrogen peroxide (Sigma-Aldrich), and a final blocking was performed with Background Sniper (Biocare Medical, Concord, CA, USA) for 10 min. Primary (FHL2 2 µg/mL and Ki67 2 µg/mL) and secondary antibodies (same as for *in vitro* immunofluorescence staining) were diluted in Dako antibody diluent (Dako, Germany). Washing steps were done in Dako washing buffer. Slides were then mounted with Vectashield antifade mounting medium (Biozol, Germany) following the manufacturer’s instructions.

Bones of the other limb of the mouse were freeze embedded following the method of the SECTION-LAB Co. Ltd. (Hiroshima, Japan). The samples were dehydrated in an ascending sucrose solution (10%, 20% and 30% in distilled water) for 24 h each at 4°C. Following this, a metal mold was placed in cooled isopropanol and filled with embedding medium (SCEM; SECTION-LAB Co. Ltd.), placing the bone in the middle. Cryosections with a thickness of 20 µm were cut following the Kawamoto method (*87*) using a cryostat (Leica CM3060S). The section was collected using a Kawamoto film (cryofilm type II C(9)) and later attached to a microscopic slide and stored at -20°C until further use. For immunostaining the slides were blocked with blocking buffer (1%BSA/ 0.1%Tween20 in PBS). Primary and secondary antibodies were diluted in blocking buffer and incubated for at least 4 and 1h at room temperature, respectively. Washing steps (2x for 15min) were conducted in washing buffer (0.1%Tween20 in PBS) and distilled water. Slides were then mounted with Dako fluorescence mounting media (S302380-2, Agilent Technologies).

### Immunofluorescence staining of human early DCCs in lymph node samples of an M0 breast cancer patient

After informed consent (ethics vote number 18-948-101), the sentinel lymph node of an M0-stage breast cancer patient (i.e. with no evidence of distant metastasis) was first divided into two halves and one half examined by histopathology. The other half was disaggregated mechanically (DAKO Medimachine, DAKO) to generate a single-cell suspension. Then, mononuclear cells were isolated using a density gradient centrifugation (60% Percoll solution, Amersham) and plated onto adhesive slides. After sedimentation of the cells, the supernatant was discarded and slides were air-dried and stored at -20°C. For immunofluorescence staining of FHL2 and Cytokeratin 8/18/19, the slides were first thawed at RT and moistened with PBS. Cells were fixed with 2% PFA (Sigma, P6148) for 10 min, washed twice with PBS and permeabilized with 0.05% TritonX100 for 5 min. After two additional washing steps with PBS, cells were blocked with PBS/10% AB serum (#805135, Bio-Rad) for 1.5 hours at RT. Primary antibody incubation with monoclonal mouse anti-human Cytokeratin 8/18/19 (2 µg/mL, clone A45-B/B3, AS Diagnostik) and polyclonal rabbit anti-human FHL2 (2 µg/mL, HPA006028, Sigma/Prestige Antibodies) was performed at 4 °C overnight in PBS/10% AB serum. Following washing with PBS (4x 20 min), the slides were incubated with secondary antibodies, anti-mouse Alexa Fluor 488 (5 µg/mL, A11029, Invitrogen) and anti-rabbit Alexa Fluor Plus 647 (5 µg/mL, A32733, Invitrogen) and DAPI (1µg/mL, MBD0015, Sigma) in PBS/5% AB serum/2.5% goat serum for 1.5 hours. After washing off the secondary antibodies (PBS 3x 10 min) the slides were mounted with SlowFade™ Glass Soft-set Antifade Mountant (S36917, Invitrogen) and imaged on Zeiss LSM980. Images were processed with ImageJ (v.1.53q) to show the max-z-projections.

### Image acquisition and analysis

Live-cell imaging was conducted in a stage top incubator (Okolab, UNO-T-H-CO_2_) mounted on an inverted epifluorescence microscope (Zeiss, AxioObserver 7) and a 10x, 0.3 NA objective at 37 °C and 5% CO_2_. Images were recorded every 6 h for 4-5 days. For FUCCI2 image analysis, live-cell imaging of cells on 2D TCP or 3D hydrogels was performed and mCherry and mVenus fluorescence signals were recorded every 6 h. Briefly, after choosing a focus plane at the beginning of the experiment for each field of view, a z-stack of this region with an interval of 20 µm between slices was acquired around this position, and phase contrast as well as fluorescence images were recorded.

At the end of the experiment, *in situ* viability staining with calcein was performed and the same z-stack/middle-focus plain as the final timelapse acquisition step was acquired. This allowed longitudinal correlation of viability at the end of the experiment with cell cycle progression over the course of the experiment. The number of cdt1-cells at the final timepoint was counted as the difference between FUCCI2+ cells (cells that either express mCherry or mVenus) at the initial and the last timepoint, with additional reference to phase contrast images. Next, viable (calcein+) single cells were counted and the dynamics of their FUCCI2 fluorescence signal extracted using a semi-automated custom-made MATLAB script. To do this, viable cells were picked and tracked over time and mean nuclear FUCCI2 intensity was extracted.

To quantify viability, projected images were thresholded for each channel corresponding to viable (calcein +) or dead (EthD-1) cells using a counting custom-made MATLAB script.

To quantify P-ERK and P-p38 intensity, the microscope Zeiss, AxioObserver 7 with a 40x 0.6 NA objective was used. Maximum projection images of single cells (n=10-20 for each condition) were manually selected based on the DAPI signal for the nucleus and phase contrast for the cell area. Next, P-ERK and P-p38 average intensity were recorded for each single cell in ZEN desk 3.8. The ratio P-ERK:P-p38 was calculated individually for each single cell.

To quantify immunofluorescence, images were first thresholded for nucleus (DAPI) and cytoskeleton (F-Actin) to localize single cells, after which the immunofluorescence signal intensity within the defined boundaries was extracted for each cell. The background signal of negatively-stained samples was used to define the threshold for positive cells. The described process was performed using an automated custom-made MATLAB script.

To quantify p21 and FHL2 localization, a Leica SP8 confocal microscope with 63x, 1.4 NA oil-immersion objective was used. Briefly, images were thresholded for nucleus (DAPI) and cytoskeleton (F-Actin) to define the respective intracellular regions, after which p21 or FHL2 average intensity within each region was recorded. The described process was performed using an automated custom-made MATLAB script.

Images of human tissue samples from patients with primary breast cancer were obtained from the Human Protein Atlas (www.proteinatlas.org) (*88*). Samples with strong FHL2 intensity were selected, of which the localization was already designated as being either nuclear or cytoplasmic.

All custom-made MATLAB scripts used in this study are available on GitHub: https://github.com/CipitriaLab/3DConfinement.

### Statistical analysis

For statistical comparison between two groups, two-tailed student’s t-test or Mann-Whitney U-test were performed for normally-distributed or non-parametric groups, respectively (*/#: p≤0.05, **/##: p≤0.01, ***/###: p≤0.001, ****/####: p≤0.0001). One-way Anova with Tukey’s correction or Kruskal-Wallis test with Dunn’s correction were conducted for multiple-group comparison. Error bars indicate mean and standard deviation used for fraction graphs. Datasets shown as box plots with median, for 25^th^ to 75^th^ percentiles and whiskers for minimum and maximum used for ratio graphs. Violin plots show dashed lines for median and dotted lines for the two quartile lines, used for fluorescence intensity graphs. GraphPad Prism software was used to plot the data and for statistical analysis.

## Acknowledgments

This work was funded by the following organizations:

- Deutsche Forschungsgemeinschaft (DFG) Emmy Noether grant (CI 203/2-1 to AC, SB, DSG, and SAEY).
- Deutsche Forschungsgemeinschaft (DFG) (KL1233/22-1 and GU 1923/3-1 to SL and CK).
- IKERBASQUE Basque Foundation for Science (AC and MMC)
- Spanish Ministry of Science and Innovation, grant PID2021-123013OB-I00 funded by MCIN/AEI/10.13039/501100011033/FEDER, UE (AC) and grant CNS2023-145020 (MMC).
- Fundación Científica Asociación Española Contra el Cáncer, grant number LABAE223466CIPI (AC)
- Osasun Saila, Eusko Jaurlaritza, grant 2023333027 (AC)
- U.H. also thanks the funding from the Basque Government Department of Education (PRE_2022_1_0006, PRE_2023_2_0096).
- International Max Planck Research School (IMPRS) on Multiscale Bio Systems (HMT).
- Helmholtz Association through program-oriented funding (JC, MG)
- Federal Ministry of Education and Research, Germany, as part of the program Health Research (BCRT; Grant No. 13GW0098) (JC, MG)
- Instituto de Salud Carlos III (ISCIII) (P21/01208), co-funded by the European Union (MMC).
- Bavarian ministry of economic affairs, energy and technology (“Aufbau einer Infrastruktur und Logistik zur translationalen Forschung mit Geweben von Krebspatienten”, AZ 20-3410.1-1-1) (CK)

The authors thank the technicians of the Research Workshop at the Charité-Universitätsmedizin Berlin for developing and manufacturing some experimental devices. We thank the lab of Dr. Joan Massagué at the Memorial Sloan Kettering Cancer Center, USA, for providing the MDA-MB-231-1833 BoM cells. We thank Aline Lueckgen, Christine Pilz, Inés Moreno-Jiménez, Geonho Song, Tina Seeman, Nicky Tam, Reynel Urrea Castellanos, Stefanie Güldener, Reinhild Dünnebacke and Laura Escobar for technical assistance with hydrogel preparation, cell culture, immunostaining, rheology, confocal microscopy, immunoblotting, blot imaging, RNA sequencing, microplate-reader measurements and cell sorting, respectively. The authors would like to acknowledge Peter Fratzl for scientific discussion.

## Author contributions

- Study design: AC, SB, UH
- Experimental work: SB, UH, SML, SMH, DSG, SAEY, AA, JC, MG
- Data analysis: SB, UH, HT, ARV, XL, SK, JW, KH, AC
- Visualization: SB, UH, ARV, AC
- Data interpretation: SB, UH, HT, ARV, SML, AA, MMC, AR, PB, SK, KH, CAK, AC
- Project supervision: AC
- Drafting manuscript: SB, UH, AC
- Revising manuscript content and approving final version: all authors

## Data and materials availability

All data needed to evaluate the conclusions in the paper are present in the paper and/or the supplementary materials. All raw and processed data, and the MATLAB codes used are available in a publicly accessible repository of the Max Planck Society (https://doi.org/10.17617/3.PWPUU0). Further, the MATLAB codes are also available in GitHub and the links are provided in the Methods section. RNAseq data have been deposited at the Gene Expression Omnibus database with accession number GSE182606.

