## Supplementary information for "Dormancy-inducing 3D-engineered matrix uncovers mechanosensitive and drug protective FHL2-p21 signaling axis"

**This PDF file includes:**

Supplementary text

Figs. S1 to S13


- Supplementary Figure S1: Norbornene-modified alginate chemical characterization
- Supplementary Figure S2: 3D mechanical confinement induces dormancy while retaining high viability
- Supplementary Figure S3: MDA-MB-231 cells within 3D Matrigel gels are resistant to doses up to 0.5 mM of 5-fluorouracil, gemcitabine and carboplatin
- Supplementary Figure S4: Position in the ranked list of genes 3D alginate stiff vs 3D Matrigel
- Supplementary Figure S5: Histogram of RNA-seq logcounts
- Supplementary Figure S6: CDK inhibitors upregulation in dormant cells compared to proliferative and independent of p53
- Supplementary Figure S7: p21 and p53 show no correlation in patients with primary estrogen receptor negative (ER-) breast cancer
- Supplementary Figure S8: p21 immunofluorescence images in 3D alginate stiff, soft and Matrigel
- Supplementary Figure S9: Position in the ranked list of genes 3D alginate soft vs stiff

- Supplementary Figure S10: p21 and FHL2 nuclear localization in 3D matrices is stiffness-dependent and suppressed upon exposure to cell proliferation regulator inhibitors

- Supplementary Figure S11: FHL2 and p21 show correlation in patients with estrogen receptor negative (ER-) breast cancer

- Supplementary Figure S12: FHL2 expression in proliferating breast cancer cell clusters in a preclinical mouse model of metastatic breast cancer

- Supplementary Figure S13: Higher FHL2 protein expression in primary breast tumor tissue and lower relapse-free survival of patients with triple-negative breast cancer and high FHL2

Tables S1 to S2

- Supplementary Table S1: Summary of the excipients or dissolvent in which drugs were prepared

- Supplementary Table S2: Forward and reverse primer sequences of dormancy genes analyzed by RT-qPCR

Captions for Movies S1 to S7

**Other Supplementary Materials for this manuscript include the following:**

Movies S1 to S7

**Supplementary Text**

a

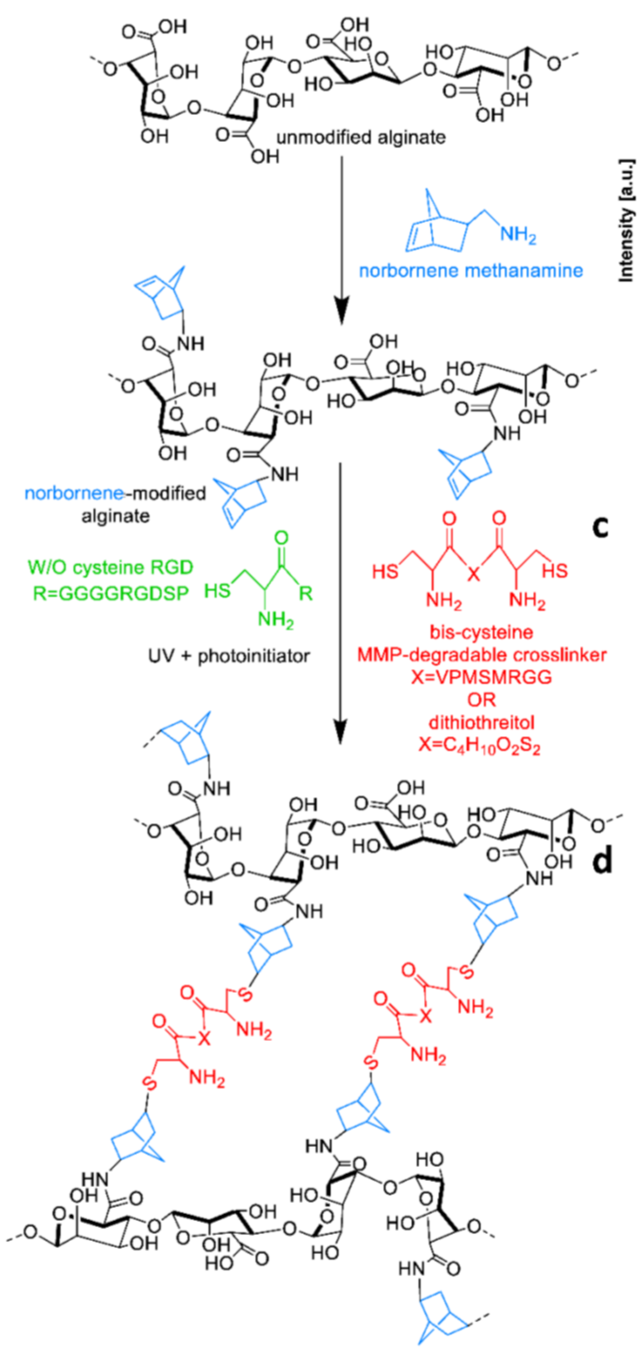


b

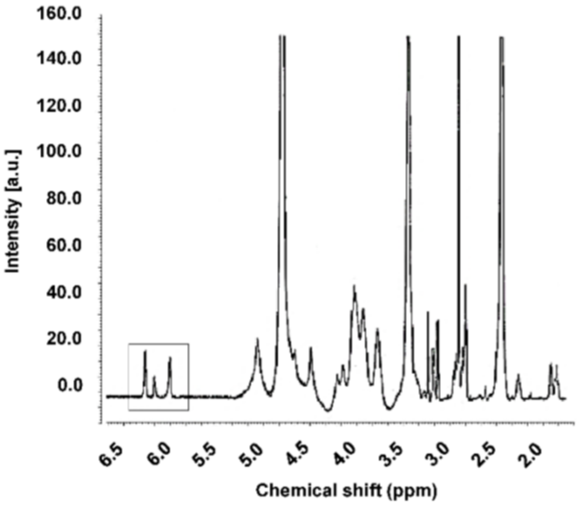


c

| | |
|---|---|
| **Molecular weight (kDa)** | 265 |
| **Norbornene DS (theo)** | 300 |
| **Norbornene DS (actual)** | 169.74 |
| **Reaction efficiency (%)** | 56.58 |

d

| | |
|---|---|
| **Sequence** | GCRD**VPMSMRGG**DRCG |
| **Structure** | $C_{63}H_{109}N_{25}O_{22}S_4$ |
| **Residues (#)** | 16 |
| **Linker** | GCRD-XXX-DRCG |
| **Molecular Weight** | 1697 |
| **Water solubility** | Good |
| **Isoelectric point** | 8.11 |
| **Charge** | +1 |

**Fig. S1 Norbornene-modified alginate chemical characterization. a)** Norbornene modification of alginate chains upon exposure to UV light and a photoinitiator in the presence of an MMP-degradable or non-degradable crosslinker via thiol-ene reaction. Cysteine-containing RGD is incorporated during the crosslinking reaction. **b)** NMR spectrum of norbornene-modified alginate, indicating the corresponding norbornene peaks (a.u.=arbitrary unit). **c)** Theoretical and actual degree of substitution (DS) with reaction efficiency (%) of norbornene-modified alginate. **d)** MMP-sensitive peptide crosslinker properties, including water solubility, isoelectric point and charge, calculated using PepCalc.com (http://pepcalc.com; Innovagen AB).

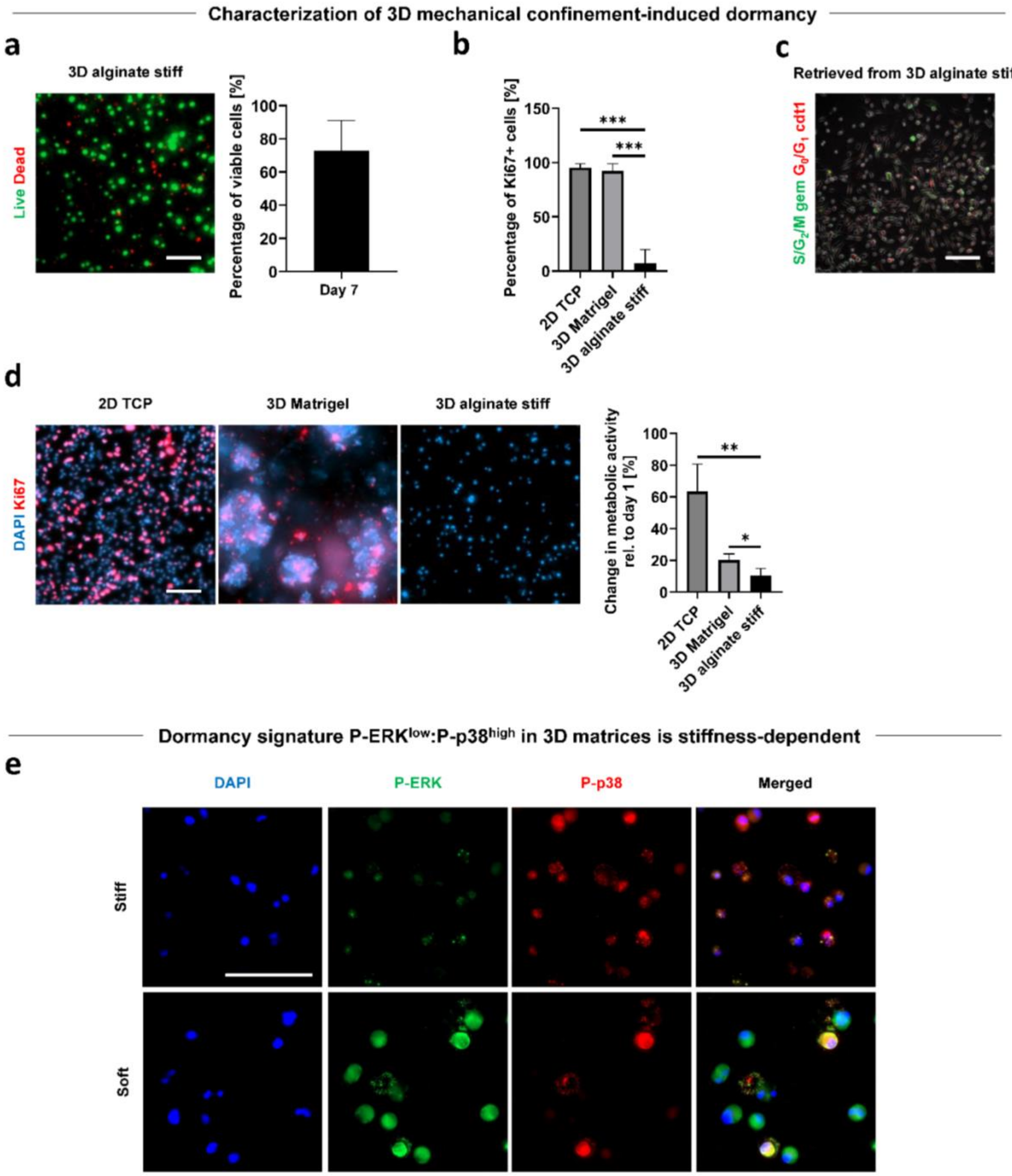


**Fig. S2 3D mechanical confinement induces dormancy while retaining high viability. a)** Representative live/dead (live=calcein, dead=ethidium homodimer) fluorescence orthogonal projection and quantification of MDA-MB-231 cells encapsulated in 3D alginate stiff after 7 days of encapsulation (n=3 gels with 134 to 204 cells). Scale bar equals 100 µm. **b)** Change in metabolic activity of MDA-MB-231 cells growing on 2D TCP, 3D Matrigel and 3D alginate stiff at day 7 relative to day 1 quantified using Presto Blue assay. **c)** MDA-MB-231-FUCCI2 cells retrieved from 3D alginate stiff after 5 days of encapsulation ($G_0/G_1$=mCherry-cdt1 and $S/G_2/M$=mVenus-geminin). Scale bar equals 100 µm. **d)** Representative Ki67 immunofluorescence images of MDA-MB-231 cells growing on 2D TCP, in 3D Matrigel and in 3D alginate stiff (blue=DAPI, red=Ki67).

Fraction of Ki67-positive (Ki67+) cells (n=3 gels for 148 to 1147 cells), student's *t*-test with respect to 3D alginate stiff, ***$p \leq 0.001$. Scale bar equals 100 μm. **e)** Representative overview maximum projection images of P-ERK and P-p38 for MDA-MB-231 cells within 3D stiff and soft alginate hydrogels (n=10-20 single cells per condition) after 5 days of encapsulation (blue=DAPI, green=P-ERK, red=P-p38). Scale bar equals 100 μm.

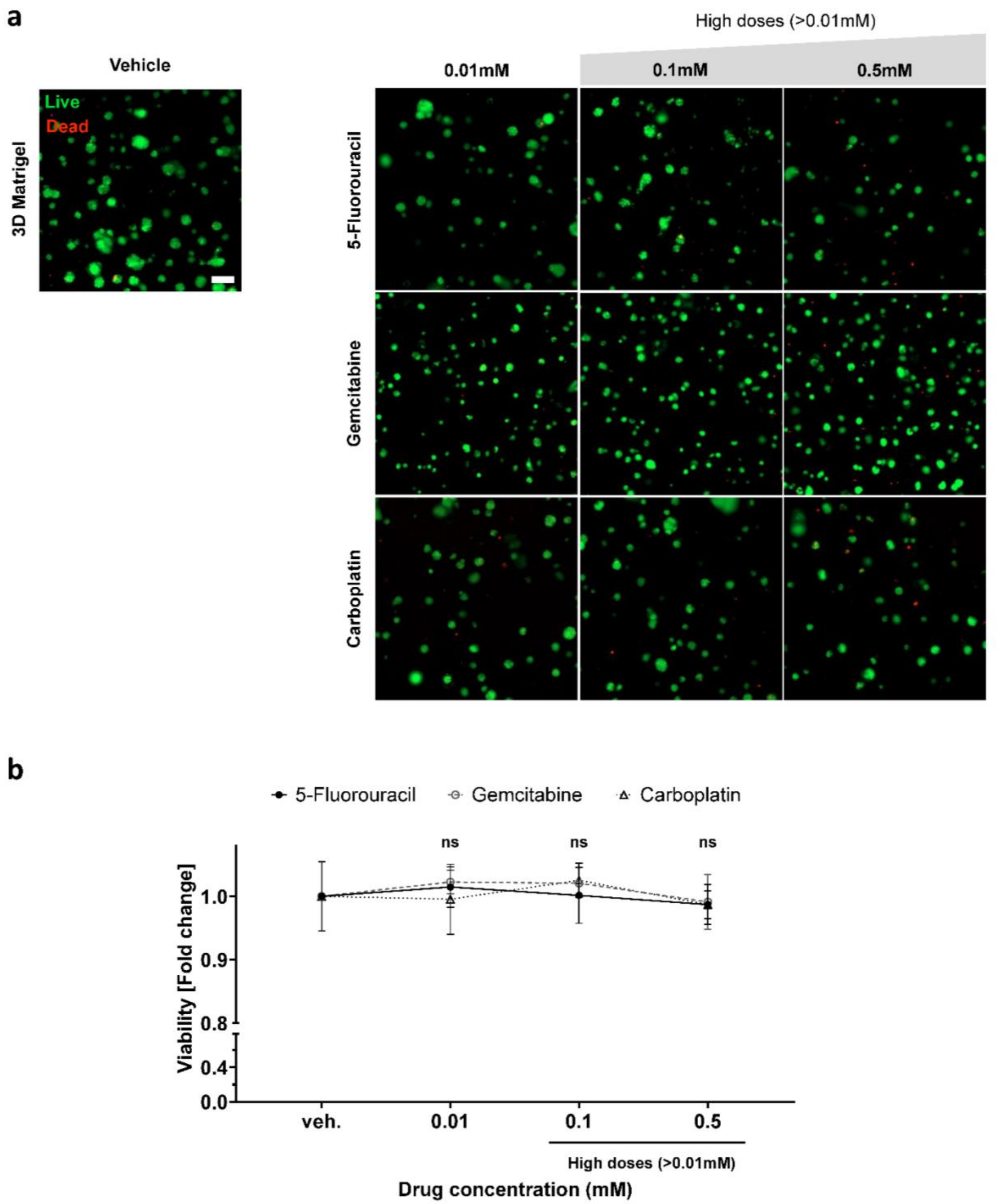


**Fig. S3 MDA-MB-231 cells within 3D Matrigel gels are resistant to doses up to 0.5 mM of 5-fluorouracil, gemcitabine and carboplatin. a)** Representative live/dead (live=calcein, dead=ethidium homodimer) maximum projection fluorescence images of MDA-MB-231 cells encapsulated in 3D Matrigel for 3 days and exposed to high doses of 5-fluorouracil, gemcitabine and carboplatin (0.01 to 0.5 mM) for 2 days (n=3 gels). Scale bar equals 100 µm. **b)** Quantification of viability fold change within 3D Matrigel with respect to the vehicle (untreated group). Statistic calculated with respect to the vehicle (untreated group). Student's t-test (n=3 hydrogels).

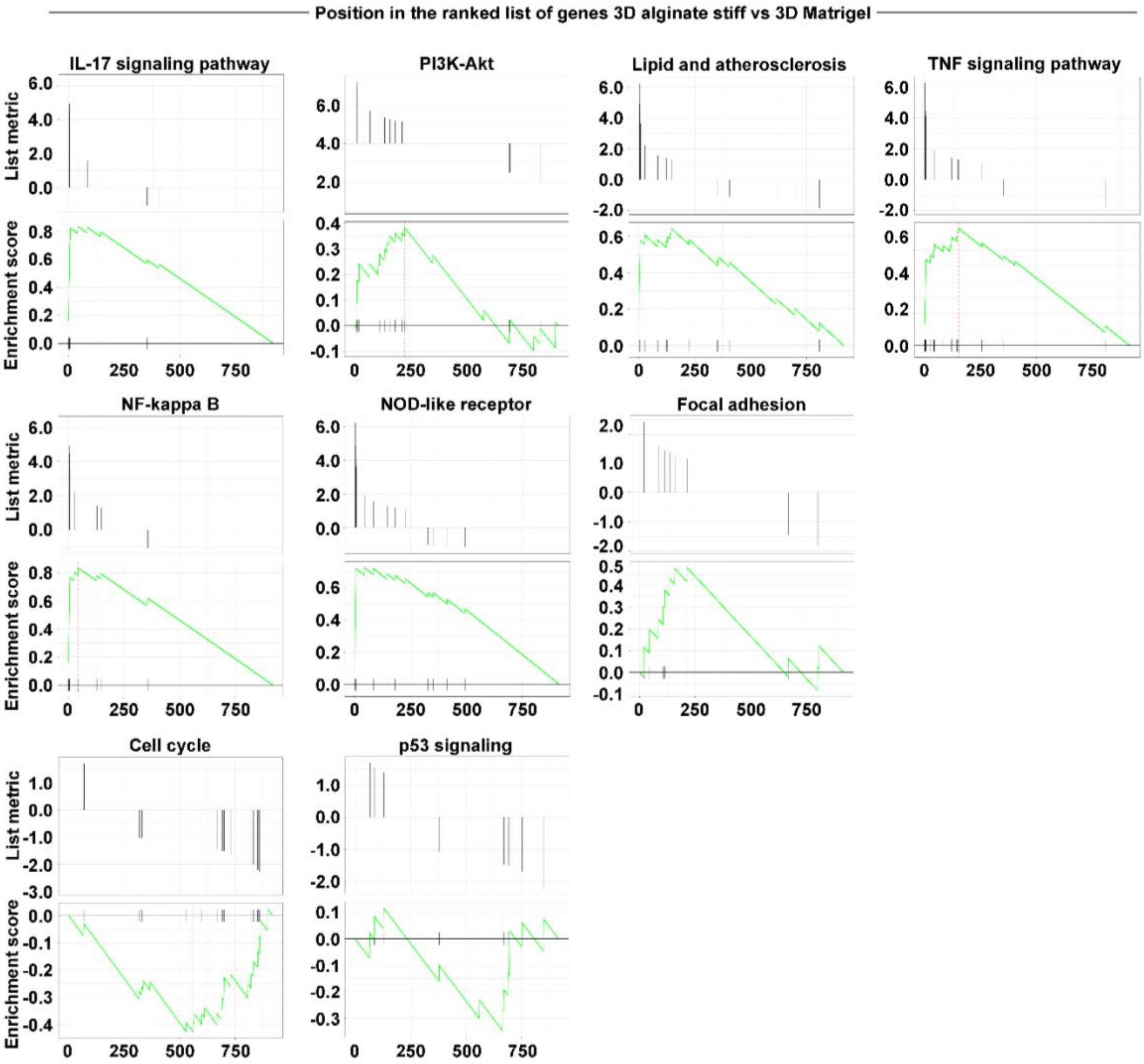


**Fig. S4 Position in the ranked list of genes 3D alginate stiff vs 3D Matrigel.** Representative genes-set-enrichment analysis (GSEA) of the differentially regulated pathways between cells grown in 3D alginate stiff and 3D Matrigel (relative to Fig.4d).

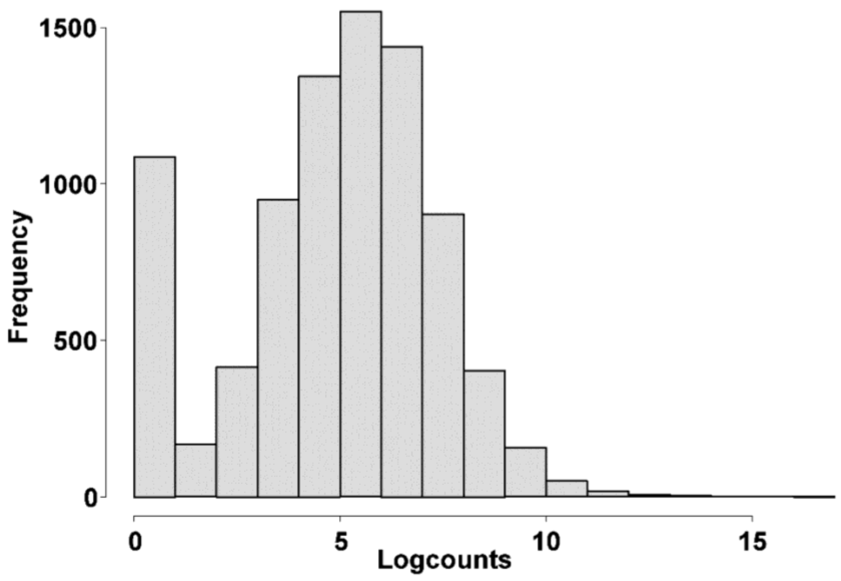


**Fig. S5 Histogram of RNA-seq log counts.**

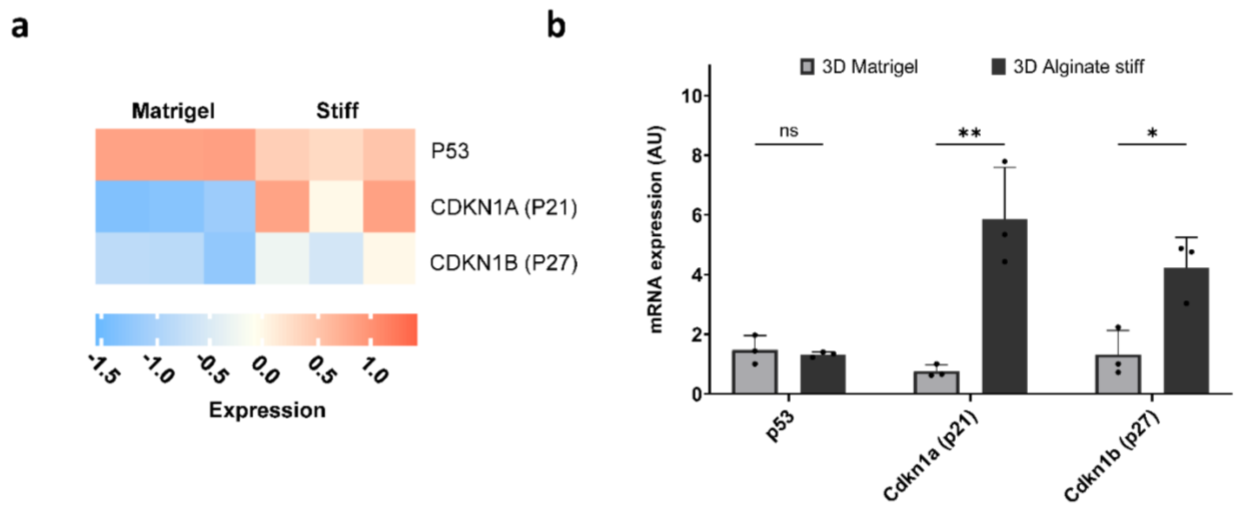


**Fig. S6 CDK inhibitors are upregulated in dormant cells compared to proliferative cells and independent of p53. a)** Heat map of differentially expressed CDK inhibitors and p53 gene obtained from RNAseq experiments of cells in 3D Matrigel vs. 3D alginate stiff. **b)** RT-qPCR validation of the mRNA expression levels of a dormancy signature for single MDA-MB-231 cells encapsulated in 3D Matrigel vs. 3D alginate stiff and retrieved 5 days later (n=3 hydrogels). Student's t-test, *$p \leq 0.05$ and **$p \leq 0.01$.

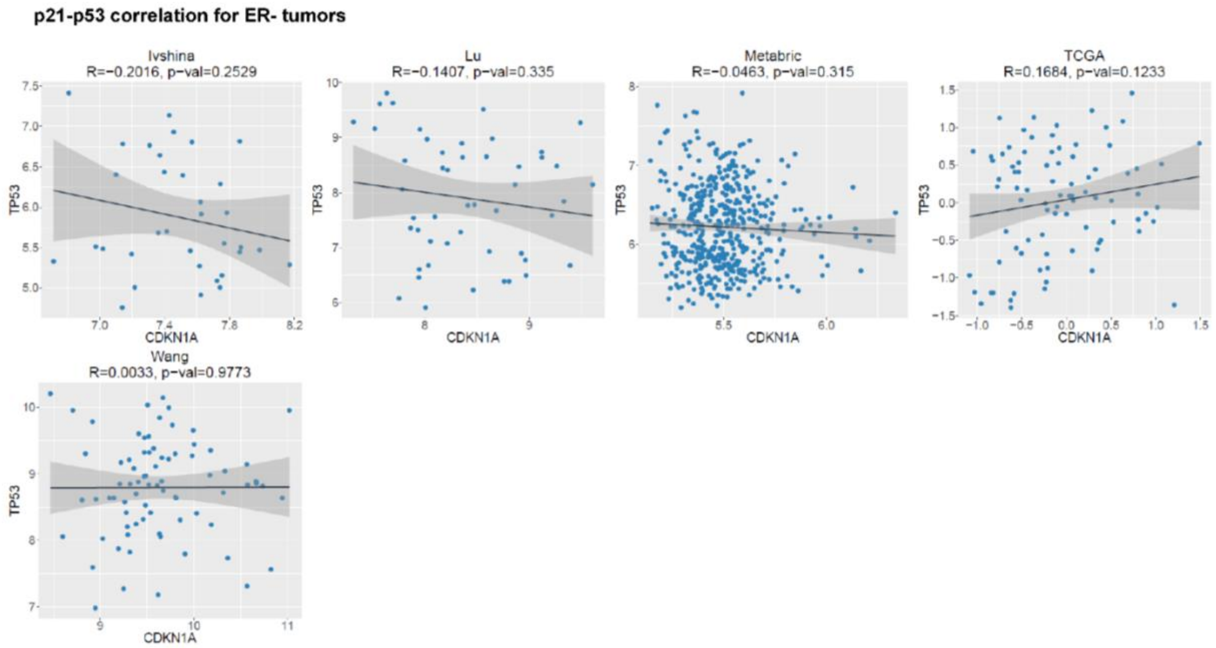


**Fig. S7 p21 and p53 mRNA levels show no correlation in patients with primary estrogen receptor negative (ER-) breast cancer.** Plotted values correspond to the log2-normalized gene expression values (fluorescence intensity or RSEM-UQ) for two genes (in X and Y-axis) for each patient in the indicated dataset. Black line represents linear regression, grey area indicates the limits of the confidence intervals and R and p indicate Pearson´s correlation coefficient (depending on the analysis selected) and statistical significance, respectively. Data obtained from CANCERTOOL[90].

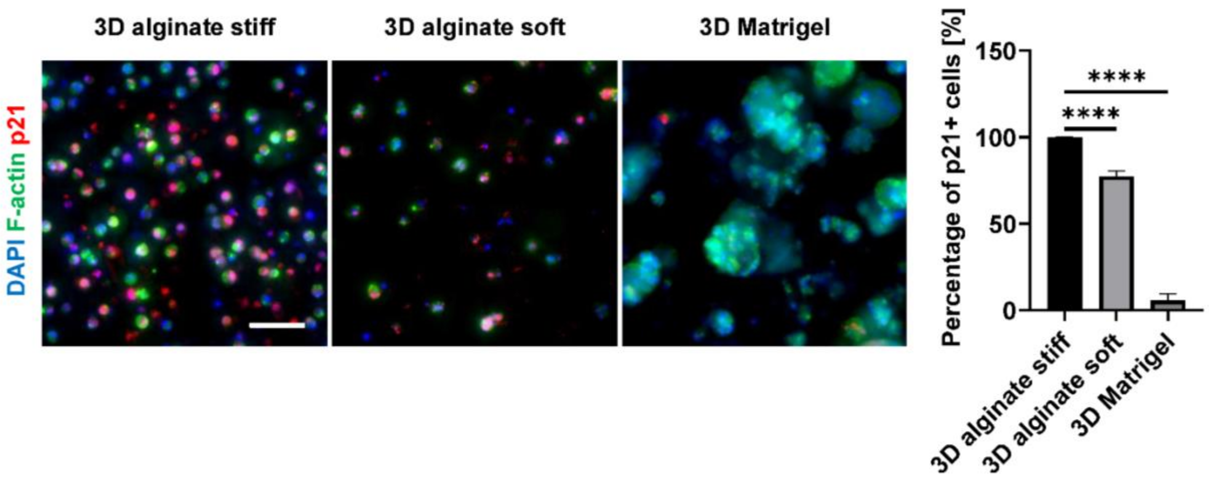


**Fig. S8 p21 immunofluorescence images in 3D alginate stiff, soft and Matrigel**. Representative images of MDA-MB-231-FUCCI2 cells in 3D alginate stiff, soft and Matrigel (blue=DAPI, green=F-Actin, red=p21). Fraction of p21-positive cells (n=4 gels for 93 to 385 cells), student's *t*-test with respect to 3D alginate stiff, ****p≤0.0001. Scale bar equals 100 µm.

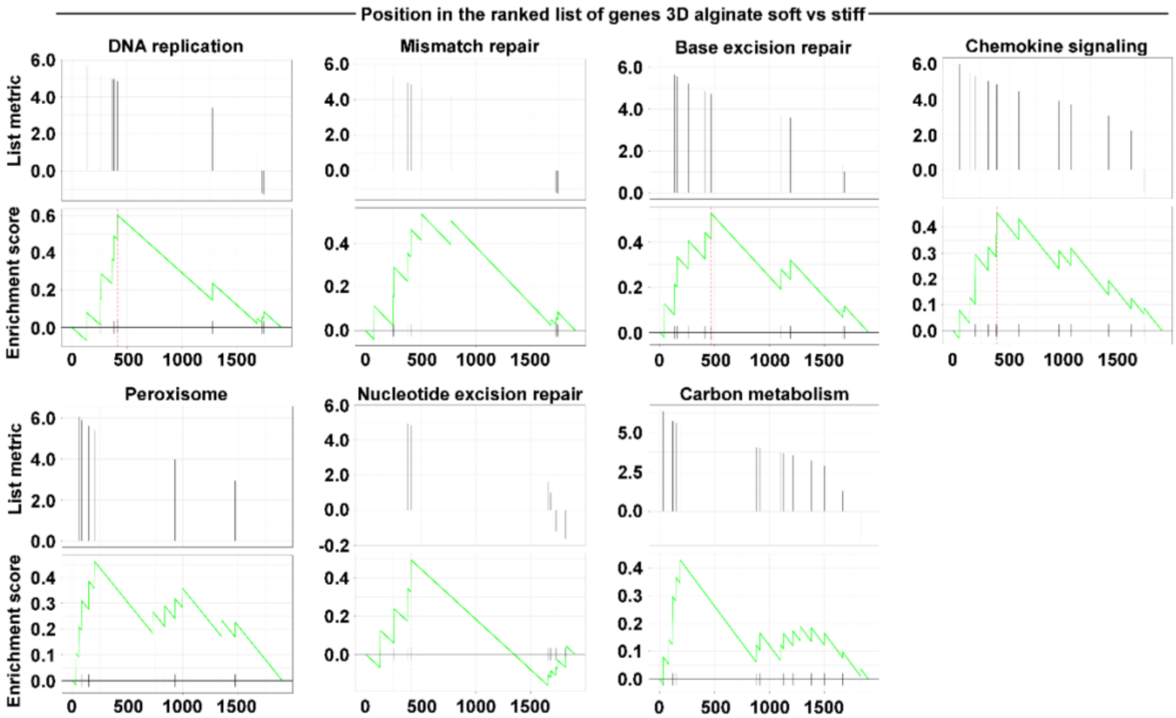


**Fig. S9 Position in the ranked list of genes 3D alginate soft vs stiff.** Genes-set-enrichment analysis (GSEA) of the differentially regulated pathways between cells grown in 3D alginate soft and stiff (relative to Fig.5a).

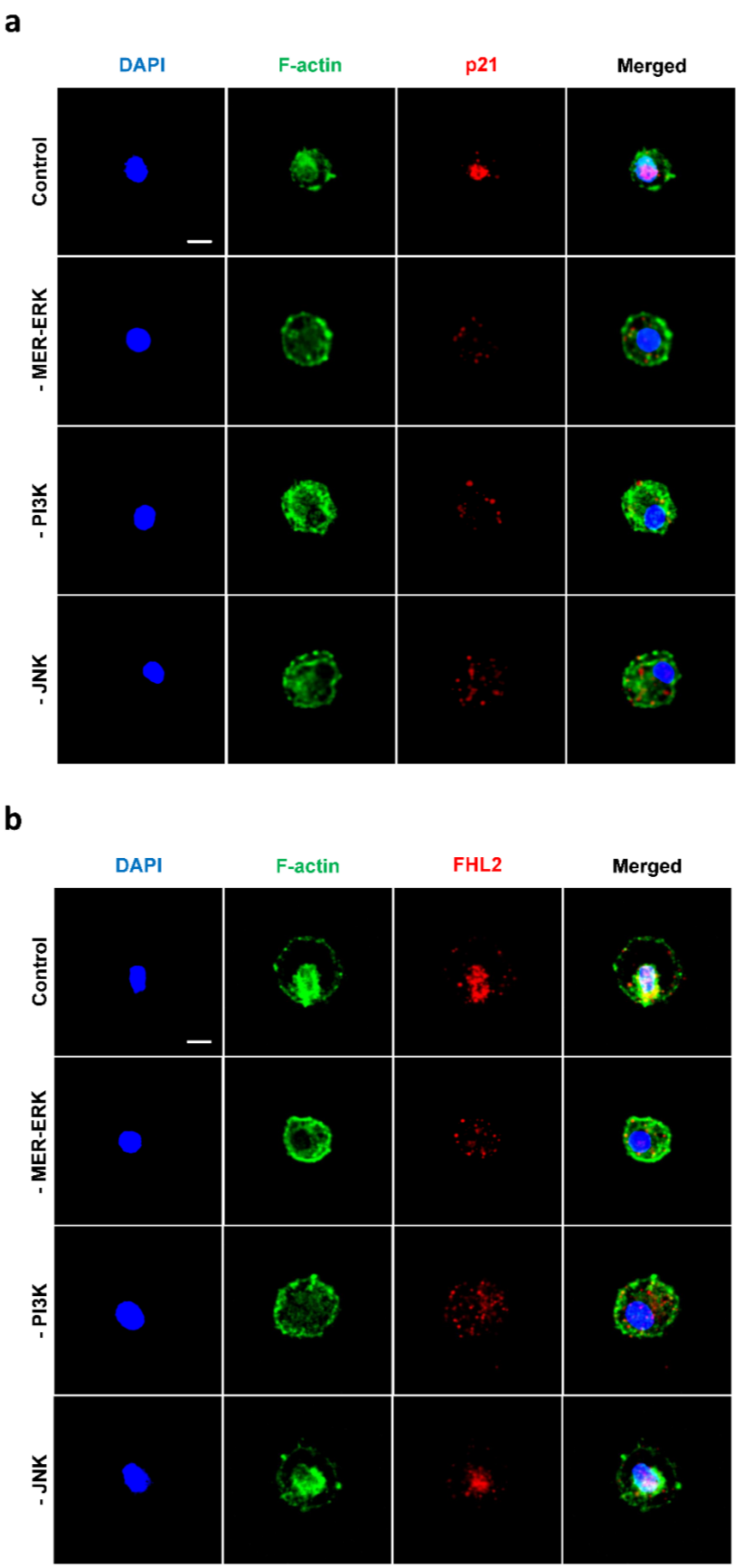
a
DAPI
F-actin
p21
Merged
Control
- MER-ERK
- PI3K
- JNK
b
DAPI
F-actin
FHL2
Merged
Control
- MER-ERK
- PI3K
- JNK

**Fig. S10. p21 and FHL2 nuclear localization in 3D matrices is stiffness-dependent and suppressed upon exposure to cell proliferation regulator inhibitors**. **a)** Representative confocal images of p21 localization for MDA-MB-231 cells within 3D stiff alginate hydrogels after 7 days in the absence or presence of indicated inhibitors (blue=DAPI, green=F-Actin, red=p21). Scale bar equals 10 μm **b)** Representative confocal images of FHL2 localization for MDA-MB-231 cells within 3D stiff alginate hydrogels after 7 days in the absence or presence of indicated inhibitors (blue=DAPI, green=F-Actin, red=FHL2). Scale bar equals 10 μm.

**FHL2-p21 correlation for ER- tumors**

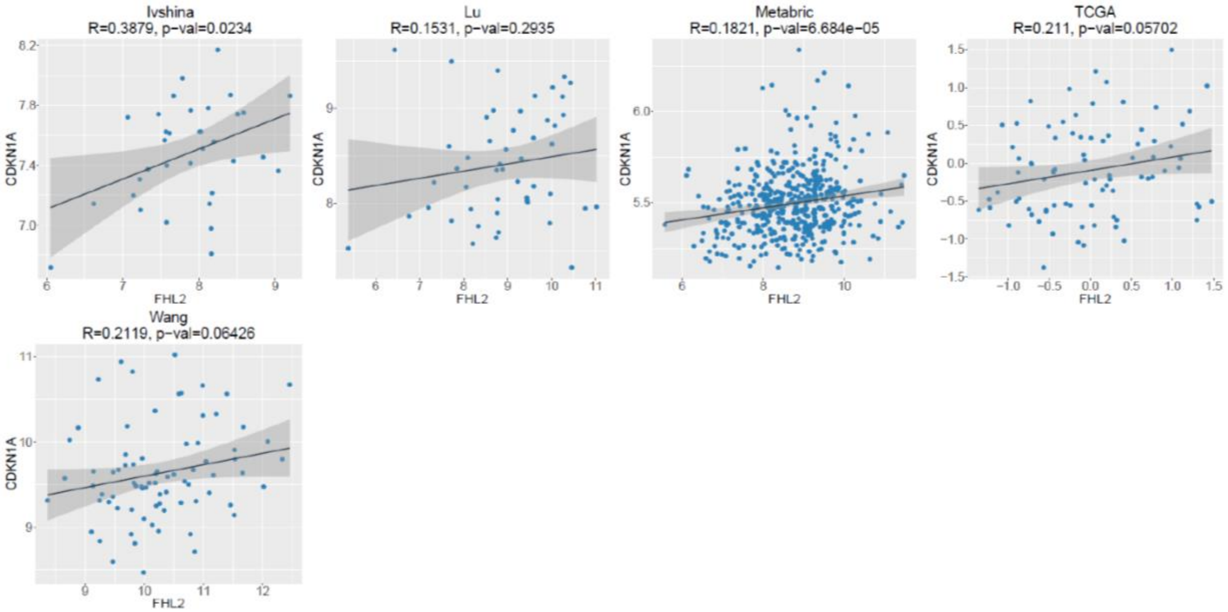


**Fig. S11 FHL2 and p21 show correlation in patients with estrogen receptor negative (ER-) breast cancer.** Plotted values correspond to the log2-normalized gene expression values (fluorescence intensity or RSEM-UQ) for two genes (in X and Y-axis) for each patient in the indicated dataset. Black line represents linear regression, grey area indicates the limits of the confidence intervals and R and p indicate Pearson´s correlation coefficient (depending on the analysis selected) and statistical significance, respectively. Data obtained from CANCERTOOL[90].

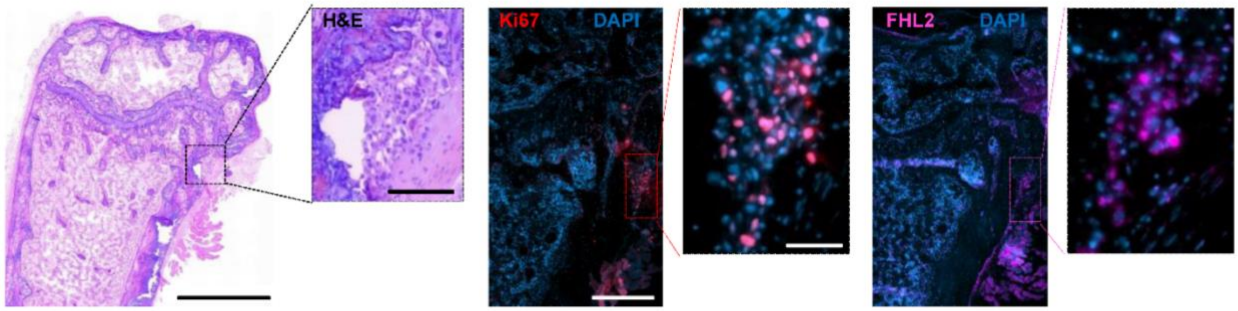


**Fig. S12 FHL2 expression in proliferating breast cancer cell clusters in a preclinical mouse model of metastatic breast cancer.** Hematoxylin and Eosin (H&E) staining of mouse femur showing an osteolytic lesion with disseminated tumor cells in the cortical region with corresponding Ki67 and FHL2 immunofluorescence staining of the selected region. Scale bar equals 1000 μm for H&E overview image, 100 μm for H&E zoomed image, 400 μm for Ki67 and FHL2 overview images and 50 μm for Ki67 and FHL2 zoomed images.

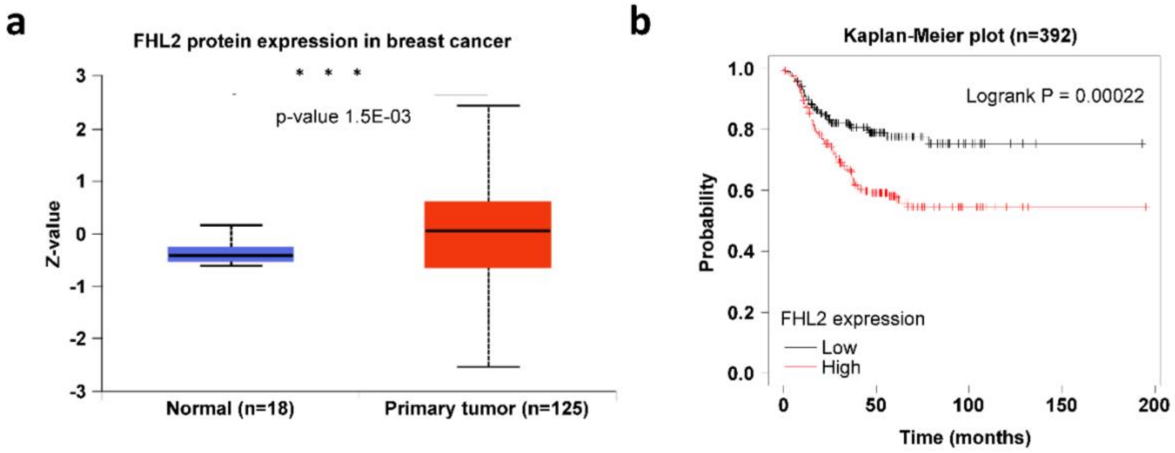


**Fig. S13 Higher FHL2 protein expression in primary breast tumor tissue and lower relapse-free survival of patients with triple-negative breast cancer and high FHL2. a)** Protein expression of FHL2 in patients with breast cancer (n=number of patients). Z-values represent standard deviations from the median across samples. Log2 Spectral count ratio values from Clinical Proteomic Tumor Analysis Consortium (CPTAC)[91] were first normalized within each sample profile, then normalized across samples. Data obtained from UALCAN database[92, 93]. **b)** Kaplan-Meier relapse-free survival plots according to FHL2 mRNA expression levels for patients with triple-negative breast cancer (n=number of patients). Data obtained from KMplot.com[94, 95].

| Drug | Excipients (Dissolvent) |
|---|---|
| Paclitaxel | Dimethyl Sulfoxide (DMSO) |
| 5-Fluorouracil | Sodium Hydroxide and Hydrochloric Acid |
| Gemcitabine | Anhydrous Disodium Phosphate, Sodium Hydroxide, Hydrochloric Acid, Anhydrous Ethanol |
| Carboplatin | - |

**Table S1. Summary of the excipients or dissolvent in which drugs were prepared.**

| Gene | Forward | Reverse |
|---|---|---|
| TP53 | CCTCCTCAGCATCTTATCCGA | GGTACAGTCAGAGCCAACCT |
| CDKN1A (p21) | TGTCTTGTACCCTTGTGCCT | AATCTGTCATGCTGGTCTGC |
| CDKN1B (p27) | ATGCGCAGGAATAAGGAAGC | TTGACGTCTTCTGAGGCCA |
| ACTB | GCAAAGACCTGTACGCCAAC | AGTACTTGCGCTCAGGAGGA |
| 18S | CGCGGTTCTATTTTGTTGGT | CGGTCCAAGAATTTCACCTC |

**Table S2. Forward and reverse primer sequences of dormancy genes analyzed by RT-qPCR**

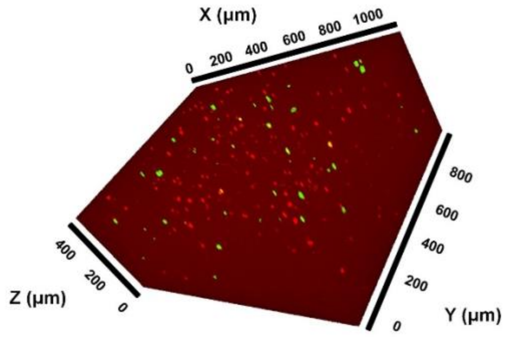


**Mov. S1 (separate file).** MDA-MB-231-FUCCI2 cells encapsulated in 3D alginate stiff after 2 days of seeding at a density of 800.000 cells/mL ($G_0/G_1$=mCherry-cdt1 and $S/G_2/M$=mVenus-geminin).

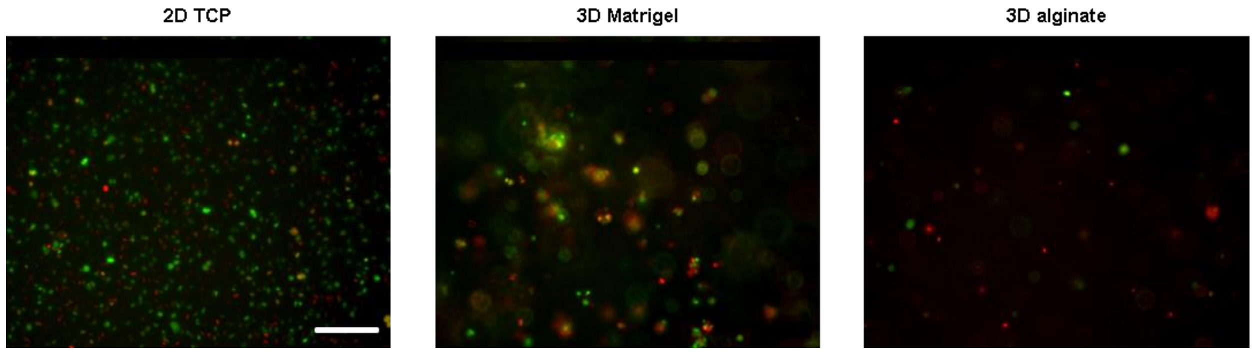


**Mov. S2-4 (separate files).** Representative time lapse videos (~4 days) of MDA-MB-231-FUCCI2 cells on 2D TCP, in 3D Matrigel and 3D alginate stiff. Scale bar equals 200 µm ($G_0/G_1$=mCherry-cdt1 and $S/G_2/M$=mVenus-geminin).

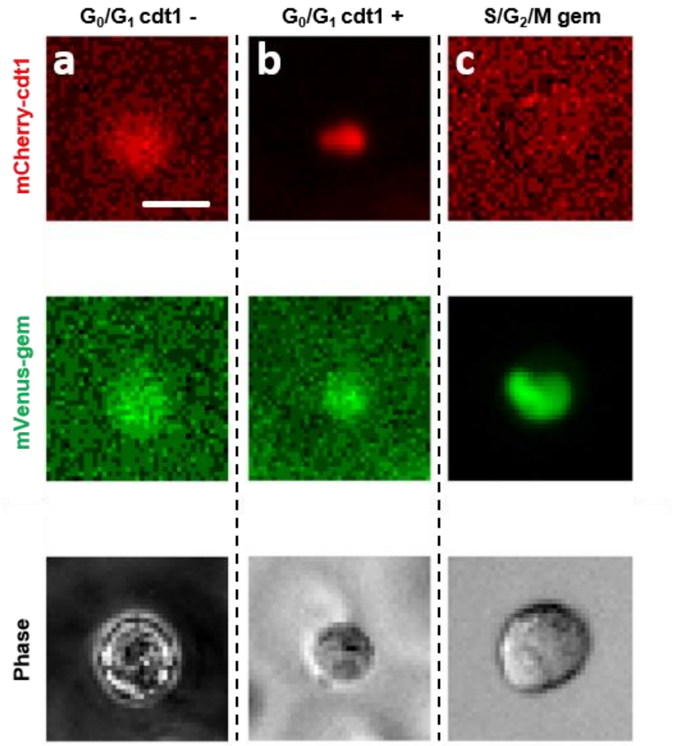


**Mov. S5-7 (separate files).** Representative time lapse videos over approximately 90 h of three separate single MDA-MB-231-FUCCI2 cells representative of a) $G_0/G_1$ cdt1-, b) $G_0/G_1$ cdt1+ and c) $S/G_2/M$ in 3D alginate stiff hydrogels ($G_0/G_1$=mCherry-cdt1, $S/G_2/M$=mVenus-geminin and phase contrast from top to bottom). The still figure shows the initial state (Day 0) of the different groups. Scale bar equals 10 μm.
